# RNA-seq preprocessing and sample size considerations for gene network inference

**DOI:** 10.1101/2023.01.02.522518

**Authors:** Gökmen Altay, Jose Zapardiel-Gonzalo, Bjoern Peters

**Author notes:** Emails : or, and, respectively.

## Abstract

**Background:** Gene network inference (GNI) methods have the potential to reveal functional relationships between different genes and their products. Most GNI algorithms have been developed for microarray gene expression datasets and their application to RNA-seq data is relatively recent. As the characteristics of RNA-seq data are different from microarray data, it is an unanswered question what preprocessing methods for RNA-seq data should be applied prior to GNI to attain optimal performance, or what the required sample size for RNA-seq data is to obtain reliable GNI estimates.

**Results:** We ran 9144 analysis of 7 different RNA-seq datasets to evaluate 300 different preprocessing combinations that include data transformations, normalizations and association estimators. We found that there was no single best performing preprocessing combination but that there were several good ones. The performance varied widely over various datasets, which emphasized the importance of choosing an appropriate preprocessing configuration before GNI. Two preprocessing combinations appeared promising in general: First, Log-2 TPM (transcript per million) with Variance-stabilizing transformation (VST) and Pearson Correlation Coefficient (PCC) association estimator. Second, raw RNA-seq count data with PCC. Along with these two, we also identified 18 other good preprocessing combinations. Any of these algorithms might perform best in different datasets. Therefore, the GNI performances of these approaches should be measured on any new dataset to select the best performing one for it. In terms of the required biological sample size of RNA-seq data, we found that between 30 to 85 samples were required to generate reliable GNI estimates.

**Conclusions:** This study provides practical recommendations on default choices for data preprocessing prior to GNI analysis of RNA-seq data to obtain optimal performance results.

## Background

Gene network inference (GNI) methods can identify putative interactions between different genes and the gene products they encode. Gene regulatory networks (GRN), which can be inferred by GNI methods, help in the basic biological understanding of genes and their functions as well as inferring potential drug targets of a disease [1]. It is worth emphasizing that the focus of this study is on GRN, which are different than co-expression networks or gene module analysis [2]. Co-expression networks identify all significant expression associations among genes, which allows clustering genes into sets (or modules) of coexpressed genes which can be further examined by e.g. Gene Set Enrichment Analysis (GSEA) [3] to associate them with biological pathways. In contrast, the goal of GRN is to infer only causal relationships between genes. An ideal GRN would connect two genes only if the gene product of one gene was directly involved in regulating the expression of the other gene, such as a transcription factor regulating the expression of another gene.

GNI methods have mostly been developed for and applied to microarray gene expression datasets [4]. The main preprocessing steps that are common to GNI algorithms include filtering, data transformation (also called scaling), normalization and estimating association scores among genes [5]. Application of GNI algorithms to RNA-seq datasets is not yet common, as RNA-seq is a relatively new technology with comparably fewer large scale datasets available [6]. RNA-seq data normalization and statistical analysis are not completely mature [7, 8] in general, and they are in an infant stage for GNI implementations. As the characteristics of RNA-seq data are different from microarray data, it may not be suitable to apply the same preprocessing methods. Unlike microarray data, distributions of RNA-seq counts are naturally heteroscedastic and larger variances are observed for larger counts [9, 10]. The skewness, mean-variance-dependency and extreme values of RNA-Seq require the use of different data preprocessing methods than the ones used for microarray [10]. For RNA-seq data, this question is well studied in the context of differential expression (DE) analysis [11–16] but not for GNI. It is thus an unanswered question how to preprocess RNA-seq data for GNI to attain best performances.

To the best of our knowledge, this is the first study that performs a systematic comparison of preprocessing combinations, which include data transformation, normalization and association estimation for GNI. We compared the combination of 15 popular preprocessing methods, along with no preprocessing case, by using 3 different GNI algorithm and 7 RNA-seq datasets for the analysis. Optimal performance was measured by comparing the identified GRN connections to a compiled literature interaction database. For this, we updated the R [17] software package, *ganet* v2.2, in order to measure the performance of each GRN prediction. Using some of the identified good performing preprocessing combinations, we conducted further analysis and determined the required sample size of RNA-seq data for optimal GNI performances. This study addresses a need of our own lab to rationale decide which pre-processing steps to apply to GNI, and we believe it provides useful guidelines and practical tools for everyone interested in infer gene networks from RNA-seq datasets.

## Results

### Analysis workflow

We assembled a set of RNA-Seq data pre-processing steps, to evaluate how they perform when applied prior to GNI analysis. We considered pre-processing steps commonly used prior to GNI analysis applied to microarray data, and combining them with pre-processing steps commonly applied to RNA-Seq data in the context of differential expression analysis. **Fig. 1** outlines the overall workflow of our analysis. Every analysis starts with RNA-Seq data as input, which provides an integer read count for each gene for each sample at **Step 1** as shown in **Fig. 1**. At **Step 2**, we consider 6 different basic data transformations: First, no conversion at all, leaving the counts as they are. Second, converting the raw counts to CPM (counts per million), which corrects for different numbers of reads in different samples. Third, we also used TPM (transcript per million) which is similar to CPM but also take gene lengths into account. For each of these three approaches, we also implemented a second version in which the transformation was followed by a logarithm in base 2 (Log-2). We refer to this group as *data types* (or *datatypes*) in the analysis and figures in the paper.

**Fig. 1.**
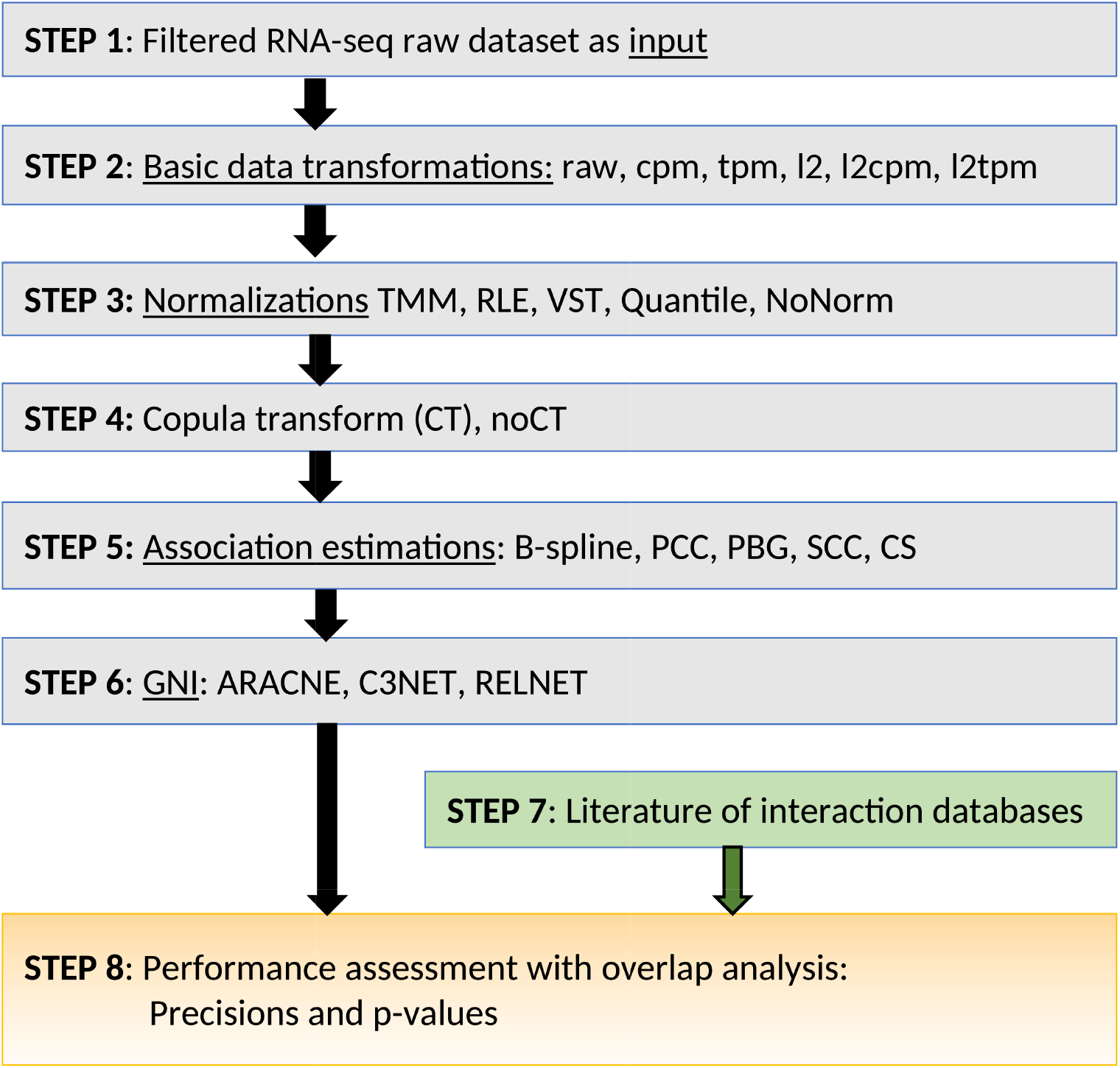
The analysis workflow for performance evaluations.

At **Step 3**, we considered more advanced normalizations, namely 3 popular normalization techniques of RNA-seq and another one that is frequently used in microarray datasets. Specifically, Variance-stabilizing transformation (VST) [18] from DeSeq2 R package [19], the trimmed mean of M-values normalization (TMM) [12] and also relative log expression (RLE) [18] from the *calcNormFactors* function of edgeR [20] R software package and finally the quantile normalization (QN) [11] that is widely used for microarray data. We also did not apply any of these 4 normalizations before GNI and named this case as no normalization (NoNorm).

**At Step 4**, we implemented the rank based data transformation technique, copula–transform (CT) as it was used in the popular GNI algorithms ARACNE [21] and C3NET [22]. They are chosen as the representative GNI algorithms along with the very base one RELNET [23]. We also did not implement CT and named this case noCT in the analysis.

**At Step 5**, we implemented association estimators to measure the similarity of the expression of gene pairs. Among the 27 available estimators [24], we selected five of the estimators that were shown to be best performing on microarray expression data [25]. They are B-spline (BS) [26], Chao-Shen (CS) [27], Pearson Correlation Coefficient (PCC), Spearman Correlation Coefficient (SCC) and Pearson-based Gaussian (PBG) as described in detail in [24, 28]. The total number of the combined preprocessing methods is 300 (6×5×2×5; see Fig. 1). Using the three GNI algorithms, we obtained 900 different performance scores for each of the 6 datasets.

**At Step 6**, we used the preprocessed datasets to infer GRNs and input them to Step 8. **At Step 7**, we input the literature interaction information to Step 8, where overlap analysis was performed between the predicted networks and the literature to evaluate the performances of each preprocessing combinations.

A pipeline was developed to implement this workflow, which is based on the workflow management system Snakemake [29].

### Exploratory data analysis

We downloaded 7 different RNA-seq datasets using the *recount* [30] R package and its repository. Details of all the used datasets are described in the Methods section. In our analysis and figures, we named them as Dataset 1 to 7 (or Data 1 to 7). In order to illustrate the effect of the preprocessing combinations on the RNA-seq data, as an example, we present the boxplots of Dataset 4 after some of the preprocessing combinations in Fig. 2. It shows how the data preprocessing methods may cause dramatic changes to a raw data. In this study, we answered the question of whether these observed changes cause improvement to the performance and if so, which specific preprocessing combinations are the best and should be used for GNI on RNA-seq datasets. From Fig. 2, we note that QN with CT seem to provide best shape from the quality control observation point of view. It allowed the median values and the shapes of boxplots over all the samples almost same to each other. In the Fig. S1 of Additional File 1, we also provided more of the other interesting RNA-seq data distributions in various forms of Dataset 2. QN produces most ordered distribution of datasets as observed in Fig. 2. Nonetheless, it does not appear to be in any of the best preprocessing combinations according to our performance evaluations. This is another point to note as an observed difference between RNA-seq and microarray, which shows that the methods developed for microarray, such as QN, may not necessarily be suitable for RNA-seq datasets too.

**Fig. 2.**
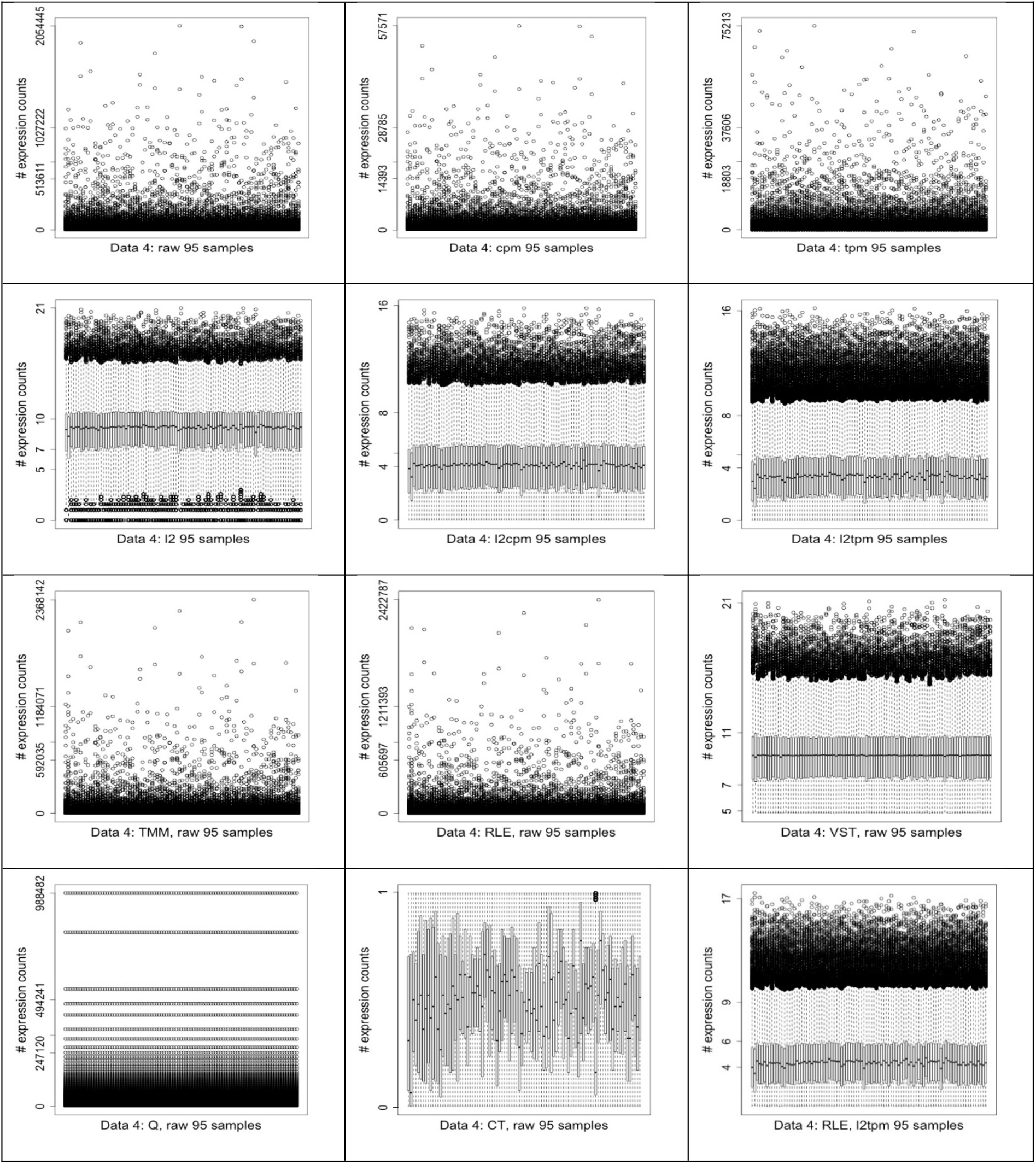
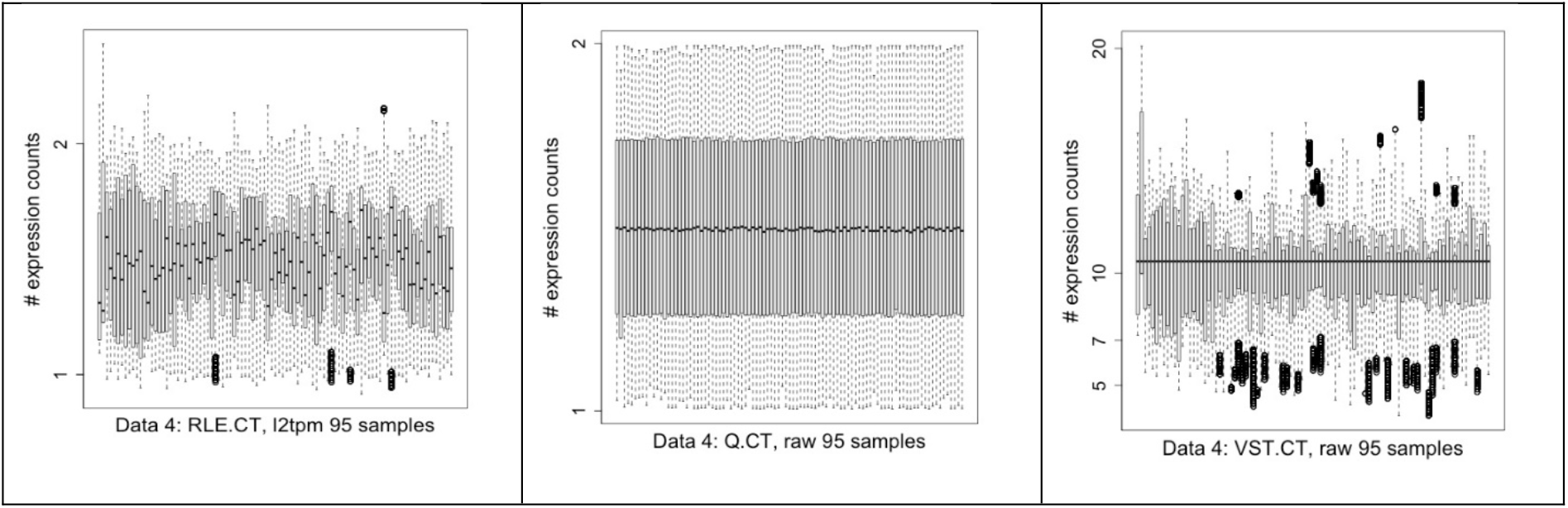
Some of the interesting RNA-seq data distributions as boxplots in various forms of Dataset 4.

### Overall performance evaluation

For the main analysis, we used 6 different RNA-seq datasets (Dataset 2 to 7) and obtained a total of 5400 different performance scores to select best performing preprocessing combinations with the representative GNI algorithms. The horizontal bar plots in Fig. 3 demonstrate 900 precision scores of all the combinations for Dataset 2 (left) and Dataset 6 (right). Similar results were provided for the other 4 datasets in Fig S2 of Additional file 1. Precisions are computed by two main components as TP (true positive), which stands for the number of correctly predicted interactions, and FP (false positive) for the number of incorrectly predicted ones. In order to determine a reference lowest point for the performance results we generated 10 random networks and assessed their precision scores. The number of interactions in the random networks selected to be the same as the one that gave best precision score over all the analysis. The mean value of the precisions of random networks computed to be approximately 0.003 and used as a reference value to compare all the results in Fig. 3. This helps to see whether the combinations result better than a random prediction. Details of generating random networks described in the Methods section.

**Fig. 3.**
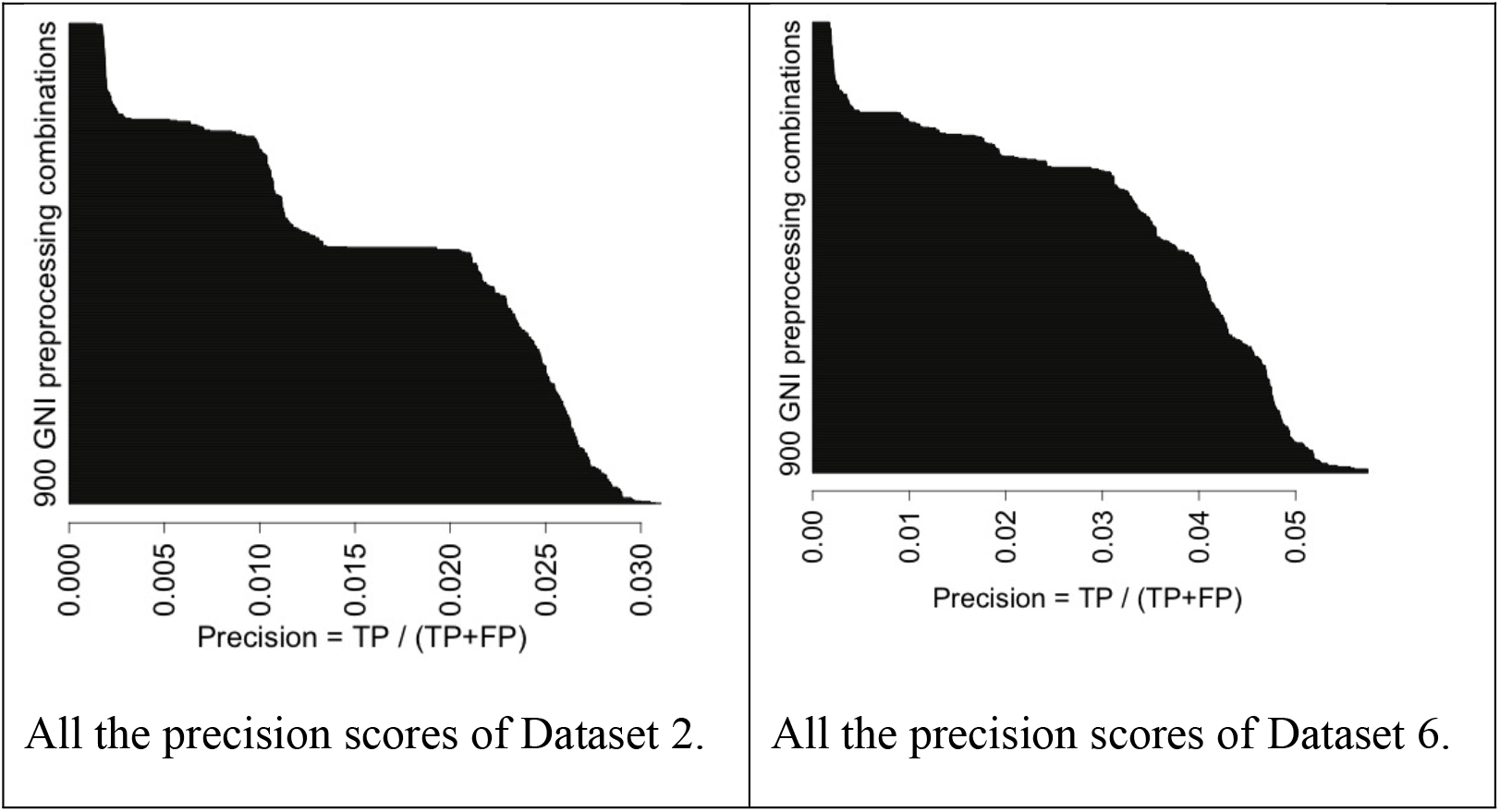
Horizontal bar plots of GNI performances over 900 different GNI and preprocessing combinations for two different datasets.

Fig. 3 appears to be quite informative. Because it shows that there are good, fair, bad and poor performing preprocessing combinations. Meaning, if one does not take careful attention on the preprocessing part of GNI applications over RNA-seq datasets, the results may be very poor similar to random predictions. CS estimator consistently provides poor results as seen in the tails of the bar plot in Fig. 3. We therefore excluded it from further analysis. On the other hand, we also observed on the right tail that a small fraction of the combinations provides good results. Among them, there several best performing ones available regarding the other good ones. When we look at the results of the other datasets (Fig S2 of Additional file 1), we observed similar trend with slightly different distributions. For example, as seen in Fig. 3, the difference between good and poor performing preprocessing combinations is sharper in the results of Dataset 2 than Dataset 6. Nonetheless, the trends of both figures are similar as all the other results, which can also be seen in Fig S2 of Additional file 1 that support our observation. This suggests that a good preprocessing combination must be carefully selected before the implementation of a GNI on RNA-seq datasets as they might influence the performances considerably.

The wide heterogeneity in the RNA-seq data makes it very difficult to choose a suitable preprocessing combination that works best for all the datasets. In Fig 4, we compare the performance of all the 6 datasets at one sight. The plot shows that each RNA-seq dataset may have dramatic effect in the performance. We excluded CS estimator for its poor performance and QN for its inconsistent performance, as will be detailed later, from this analysis. Effects of each algorithm were also evaluated separately in the coming sections.

**Fig. 4.**
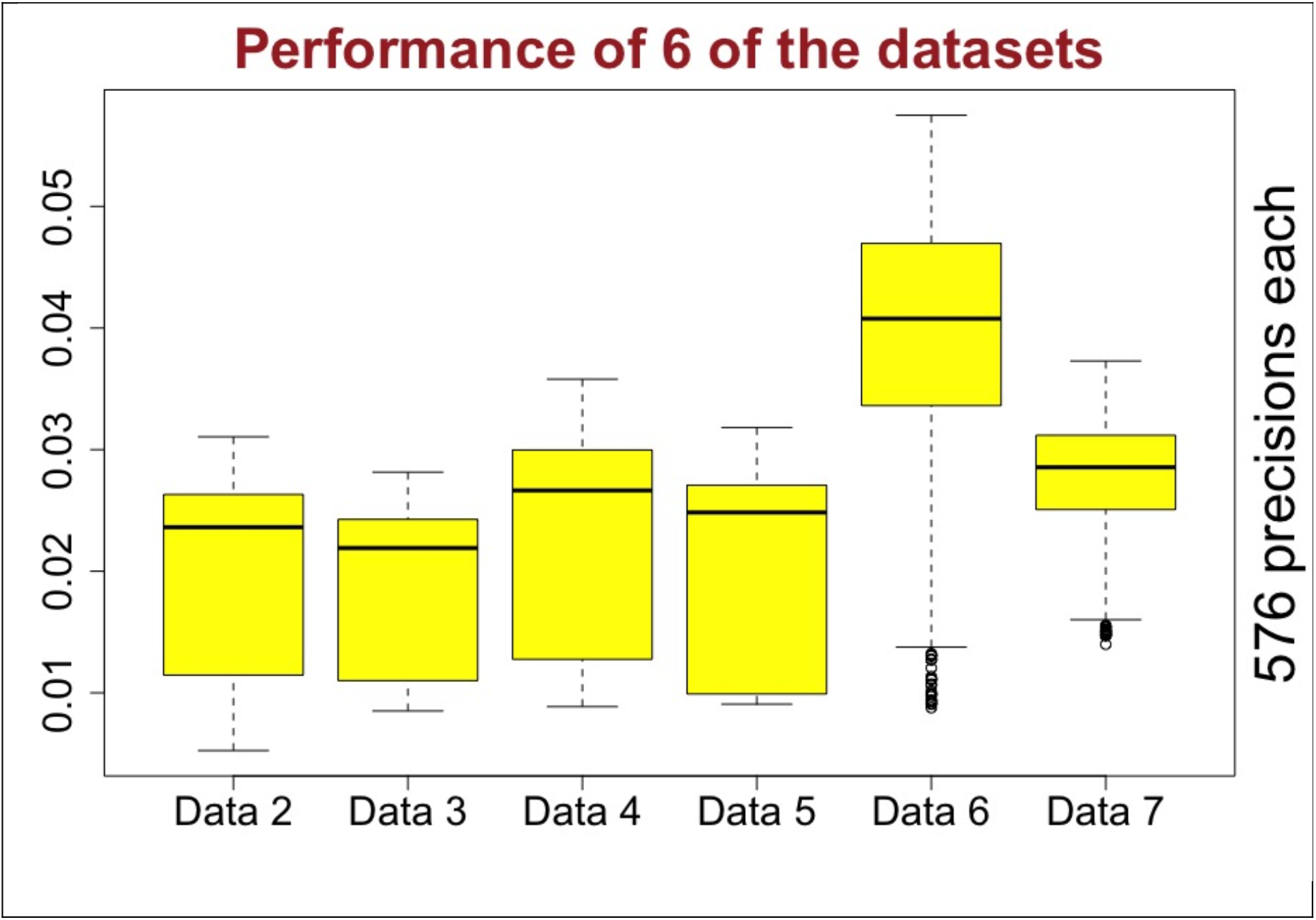
Performances of 6 of the datasets for various preprocessing and GNI combinations.

In Table 1, we zoom into the results of Dataset 6 presented in Fig.3. It shows the top 20 best performing GNI and preprocessing combinations with respect to precision scores. Table 1 also shows, in the last three row, the more statistically significant (lower p-value) overlapping analysis results but with poor precision scores as a noteworthy example. In Table 1, we presented the highest precision scores comparing 5400 results of all the 6 datasets. It stands as an exemplary table that shows the performance results of our analysis in detail. We derived all the figures for the analysis from those information of all the datasets.

**Table 1.** Shows some of the inference performances of the top 20 and the 3 very low p-value preprocessing combinations of Dataset 6.

| Combinations | Precision<br>TP/(TP+FP) | #Overlap<br>(TP) | #Predicted<br>(TP+FP) | p.value |
| --- | --- | --- | --- | --- |
| c3.PBG_CT_NoNorm_l2tpm | 0.0575 | 273 | 4750 | 4.75E-194 |
| c3.PBG_CT_NoNorm_tpm | 0.0575 | 273 | 4750 | 4.75E-194 |
| c3.PCC_CT_NoNorm_l2tpm | 0.0575 | 273 | 4750 | 4.75E-194 |
| c3.PCC_CT_NoNorm_tpm | 0.0575 | 273 | 4750 | 4.75E-194 |
| c3.SCC_CT_NoNorm_l2tpm | 0.0575 | 273 | 4750 | 4.75E-194 |
| c3.SCC_CT_NoNorm_tpm | 0.0575 | 273 | 4750 | 4.75E-194 |
| c3.SCC_noCT_NoNorm_l2tpm | 0.0575 | 273 | 4750 | 5.59E-194 |
| c3.SCC_noCT_NoNorm_tpm | 0.0575 | 273 | 4750 | 5.59E-194 |
| c3.bspline_CT_NoNorm_l2tpm | 0.0561 | 270 | 4820 | 3.72E-189 |
| c3.bspline_CT_NoNorm_tpm | 0.0561 | 270 | 4820 | 3.72E-189 |
| c3.PBG_noCT_NoNorm_l2tpm | 0.0557 | 264 | 4740 | 3.22E-184 |
| c3.PCC_noCT_NoNorm_l2tpm | 0.0557 | 264 | 4740 | 3.22E-184 |
| c3.PBG_noCT_NoNorm_l2cpm | 0.0544 | 263 | 4840 | 5.57E-181 |
| c3.PCC_noCT_NoNorm_l2cpm | 0.0544 | 263 | 4840 | 5.57E-181 |
| c3.bspline_noCT_RLE_l2 | 0.0542 | 263 | 4850 | 1.20E-180 |
| c3.bspline_noCT_NoNorm_l2 | 0.0534 | 255 | 4780 | 1.42E-173 |
| c3.bspline_noCT_TMM_l2 | 0.0533 | 259 | 4860 | 3.20E-176 |
| ar.PBG_noCT_NoNorm_l2tpm | 0.0533 | 415 | 7790 | 7.20E-281 |
| ar.PCC_noCT_NoNorm_l2tpm | 0.0533 | 415 | 7790 | 7.20E-281 |
| c3.bspline_noCT_TMM_l2cpm | 0.0527 | 263 | 4990 | 1.36E-177 |
| ..... | ..... | ..... | ..... | ..... |
| rn.PCC_CT_VST_l2cpm | 0.00936 | 5300 | 566000 | 0 |
| rn.SCC_noCT_VST_l2tpm | 0.00908 | 5150 | 567000 | 0 |
| rn.bspline_noCT_VST_l2tpm | 0.00878 | 4960 | 565000 | 0 |

There are shortened representations of the preprocessing methods in the first column of Table 1. Most of them are available in the main text; others expressed here: ar, c3, and rn are ARACNE, C3NET and RELNET GNI algorithms, respectively. If a name is not mentioned it means it was not used but some of them emphasized with ‘No’ prefix to mention non-existence. Namely, noCT and NoNorm means copula transform (CT) and normalizations are not used, respectively. If there is no Log-2 applied, then there is no l2 in the names. Overlap analysis over the literature via hyper-geometric tests resulted the p-values. #Overlap represents the number of overlap between the predicted gene network and the literature interactions database.

If we had looked at a result of a single dataset we would end up selecting a best preprocessing combination. But, we used 6 different datasets that allowed us to find out that there is no best preprocessing combination valid for all the datasets. Rather, there are good combinations that work well mostly but each dataset might need a different preprocessing combination for the best GNI performance. This is pretty different than we would expect to find in the beginning of this study. For example, the best performing GNI and preprocessing combination in Table 1 does not provide highest scores in the other datasets. Therefore, in order to be able to select the good performing preprocessing combinations, we first evaluated them for each GNI algorithm separately to eliminate the effect of GNI and see only the performances of preprocessing combinations. We then ranked them based on precision scores for each dataset and obtained 6 different rank tables; each have ranked values for each unique preprocessing combinations. We computed the median values of these ranks over all the 6 datasets. We then sorted all the combinations based on the median ranks. This approach allowed us to see the performance of each preprocessing combinations considering all the 6 datasets at once. If there is a tie, then we assigned the minimum rank to each of them. Best rank is 1 and worst rank is 192 as we excluded some of the poor performing methods (CS and QN) already from the analysis. We present the top 25 best performing results from this analysis in Table 2 and provide all of them in Additional file 2.

**Table 2.** Top 25 ranked GNI performances with ARACNE over all the 6 datasets. Sorted by median ranks. The worst rank can be 192 and the best can be 1.

| GNI preprocessing combinations | median rank | mean rank | best rank | worst rank |
| --- | --- | --- | --- | --- |
| ar.bspline_CT_NoNorm_l2tpm | 22.5 | 57.2 | 7 | 143 |
| ar.bspline_CT_NoNorm_tpm | 22.5 | 57.2 | 7 | 143 |
| ar.PBG_noCT_NoNorm_tpm | 24.5 | 48.5 | 3 | 165 |
| ar.PCC_noCT_NoNorm_tpm | 24.5 | 48.5 | 3 | 165 |
| ar.PBG_CT_NoNorm_l2tpm | 31.5 | 61.3 | 9 | 156 |
| ar.PBG_CT_NoNorm_tpm | 31.5 | 61.3 | 9 | 156 |
| ar.PCC_CT_NoNorm_l2tpm | 31.5 | 61.3 | 9 | 156 |
| ar.PCC_CT_NoNorm_tpm | 31.5 | 61.3 | 9 | 156 |
| ar.PBG_noCT_NoNorm_l2tpm | 33.5 | 47.3 | 1 | 154 |
| ar.PCC_noCT_NoNorm_l2tpm | 33.5 | 47.3 | 1 | 154 |
| ar.SCC_CT_NoNorm_l2tpm | 33.5 | 63 | 8 | 160 |
| ar.SCC_CT_NoNorm_tpm | 33.5 | 63 | 8 | 160 |
| ar.bspline_noCT_TMM_l2 | 37 | 72.8 | 5 | 187 |
| ar.bspline_noCT_RLE_l2 | 38.5 | 75.3 | 6 | 188 |
| ar.SCC_noCT_NoNorm_l2tpm | 40 | 66.5 | 6 | 162 |
| ar.SCC_noCT_NoNorm_tpm | 40 | 66.5 | 6 | 162 |
| ar.bspline_noCT_RLE_raw | 40.5 | 80.2 | 21 | 176 |
| ar.PBG_noCT_TMM_l2tpm | 40.5 | 57.2 | 1 | 175 |
| ar.PCC_noCT_TMM_l2tpm | 40.5 | 57.2 | 1 | 175 |
| ar.bspline_noCT_NoNorm_l2tpm | 41 | 76.5 | 1 | 190 |
| ar.PBG_CT_TMM_l2tpm | 42 | 74 | 7 | 171 |
| ar.PCC_CT_TMM_l2tpm | 42 | 74 | 7 | 171 |
| ar.SCC_CT_TMM_l2tpm | 42 | 75.3 | 9 | 173 |
| ar.SCC_noCT_TMM_l2tpm | 42 | 75.3 | 9 | 173 |
| ar.bspline_noCT_NoNorm_l2 | 44.5 | 68.3 | 17 | 174 |

The complete tables for each GNI algorithm and the table of the similar analysis of all the GNI and preprocessing combinations together can be seen as a spreadsheet file in Additional file 2 that allows sorting the results as wanted. Table 2 is very useful to see that there is no best performing preprocessing combination in general. For example, the top-ranking prepressing combination of ARACNE GNI algorithm is the combination of B-spline association estimator, CT, no normalization and Log-2 TPM data. There is also a tie and the same combination with TPM instead of Log-2 TPM performs the same. But even the best combination has median rank = 22.5, best rank = 7 and worst rank = 143 out of 192. This shows that even the best performing combination may still provide very poor results. This is a very important observation out of this study, which cause us to be always very cautious in GNI of RNA-seq data. We have a conclusion and suggest a solution based on this observation in the Conclusion section.

The preprocessing combinations in Table 2 may be considered to be in the set of good performing ones for ARACNE GNI algorithm with some cautious. We present similar analysis for C3NET GNI algorithm in Table 3. As seen, the top performing preprocessing combinations are very similar to the results of ARACNE with some difference in their ranks. The best performing combinations for C3NET is B-spline, CT and TPM (or Log-2 of TPM) has median rank of 10.5, best rank 2 and worst rank 148. Again, we see that the best performing combinations has still potential to perform very poorly.

**Table 3.** Top 25 ranked GNI performances with C3NET over all the 6 datasets. Sorted by median ranks and the worst rank can be 192 and the best can be 1.

| GNI preprocessing combinations | median rank | mean rank | best rank | worst rank |
| --- | --- | --- | --- | --- |
| c3.bspline_CT_NoNorm_l2tpm | 10.5 | 47.7 | 2 | 148 |
| c3.bspline_CT_NoNorm_tpm | 10.5 | 47.7 | 2 | 148 |
| c3.PBG_noCT_VST_l2tpm | 18.5 | 54.7 | 1 | 190 |
| c3.PCC_noCT_VST_l2tpm | 18.5 | 54.7 | 1 | 190 |
| c3.bspline_noCT_VST_l2tpm | 20.5 | 47 | 4 | 192 |
| c3.bspline_noCT_TMM_l2 | 23 | 70.5 | 7 | 183 |
| c3.PBG_noCT_NoNorm_l2tpm | 26 | 57.3 | 5 | 170 |
| c3.PCC_noCT_NoNorm_l2tpm | 26 | 57.3 | 5 | 170 |
| c3.bspline_noCT_RLE_l2 | 29 | 70.3 | 3 | 186 |
| c3.bspline_noCT_NoNorm_tpm | 31.5 | 69.8 | 6 | 172 |
| c3.PBG_CT_NoNorm_l2tpm | 41 | 66.3 | 1 | 180 |
| c3.PBG_CT_NoNorm_tpm | 41 | 66.3 | 1 | 180 |
| c3.PCC_CT_NoNorm_l2tpm | 41 | 66.3 | 1 | 180 |
| c3.PCC_CT_NoNorm_tpm | 41 | 66.3 | 1 | 180 |
| c3.PBG_CT_VST_l2tpm | 45 | 64.3 | 5 | 174 |
| c3.PCC_CT_VST_l2tpm | 45 | 64.3 | 5 | 174 |
| c3.SCC_CT_VST_l2tpm | 45 | 62.8 | 5 | 174 |
| c3.SCC_noCT_VST_l2tpm | 45 | 62.8 | 5 | 174 |
| c3.SCC_CT_NoNorm_l2tpm | 46 | 68 | 1 | 180 |
| c3.SCC_CT_NoNorm_tpm | 46 | 68 | 1 | 180 |
| c3.bspline_noCT_VST_l2 | 47 | 68.7 | 25 | 150 |
| c3.PBG_noCT_RLE_l2tpm | 47.5 | 70.5 | 7 | 159 |
| c3.PCC_noCT_RLE_l2tpm | 47.5 | 70.5 | 7 | 159 |
| c3.SCC_noCT_NoNorm_l2tpm | 48 | 72.8 | 7 | 186 |
| c3.SCC_noCT_NoNorm_tpm | 48 | 72.8 | 7 | 186 |

Ranking approach, as in Table 2 and 3, helps to evaluate the combinations over all the 6 datasets at once but we still need to consider the raw information as in Table 1. Because there are many good performing combinations with very close precision scores. For example, C3NET precision scores of Dataset 2 ranges from 0.024 to 0.03 for all the 192 combinations (Additional file 2, sheet 3). Also, the top 53 combinations of it is ranging from 0.028 to 0.0308. In this list of results, we had removed the poor performing method CS and QN. If we look at all the results in all the 6 datasets for C3NET and ARACNE, as shown in Fig. 5, the performances are close to each other in 5 out of 6 datasets but for one of them it is considerably higher. This shows that performances of the good combinations are quite close to each other mostly when the very poor performing ones are not used.

**Fig. 5.**
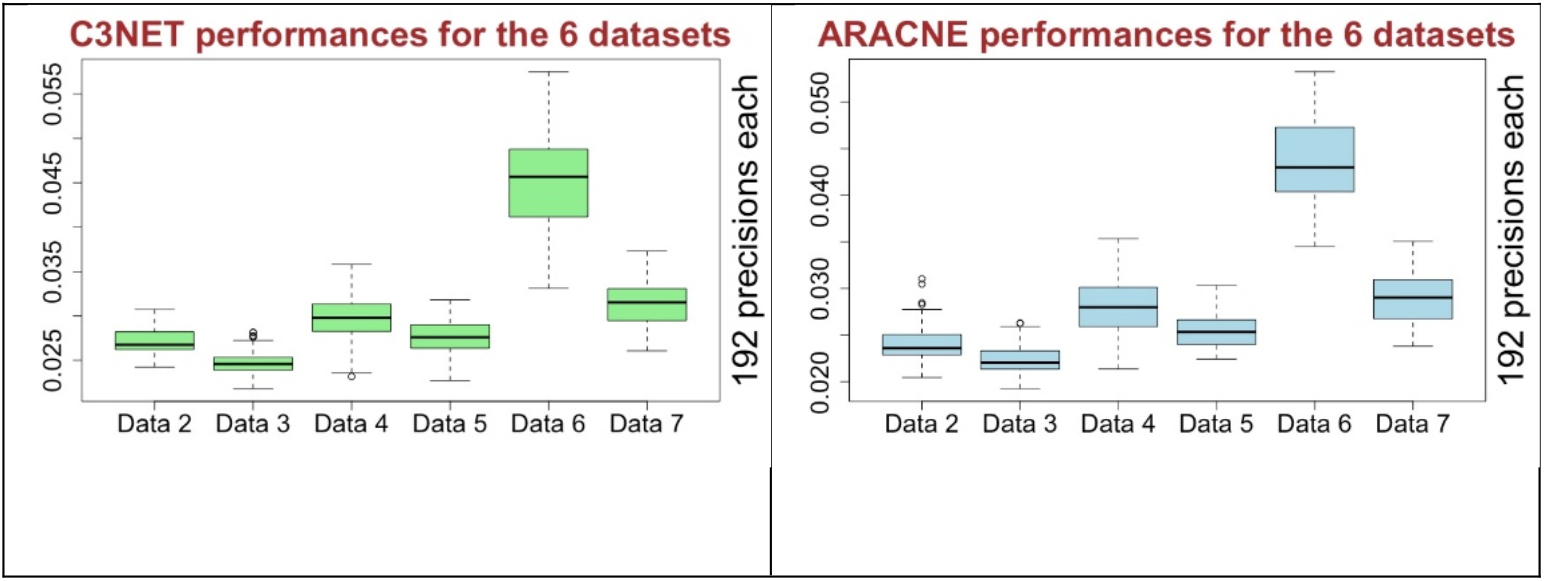
Performance of C3NET and ARACNE over all datasets (CS and QN excluded).

On the other hand, there are still best performing combinations available and that small difference in accurate predictions may still biologically have profound effect. Because even a single newly detected interaction may be the main regulator of that biological condition. Therefore, it is still interesting to use the best performing ones among the good ones when inferring GRN from RNA-seq data. In order to be able to find best ones, we use the ranking approach on them as explained above. The difference among rank values may look larger than the real performance scores. Therefore, we consider all the information as in Table 1, 2 and 3 (Additional file 2) while selecting a good combination. This approach lead us to determine the good preprocessing combinations as follows in the quotes: ‘B-spline and CT and Log-2 TPM (or TPM)’, ‘PCC or PBG and no CT and VST and Log-2 TPM’, ‘B-spline and no CT and VST and Log-2 TPM’. RLE and TMM may also be replaced in the estimators of those combinations. As observed, even the selected good ones may not always provide best performances. If the best performing one is wished to be selected for a specific dataset, then all the determined good ones should be tested along with the several other good preprocessing combinations mentioned earlier on Table 2. In the following, we analyze each step of preprocessing, shown in Fig. 1, separately. This will help us to learn the individual effects of the methods for each step in general.

### Effect of transformed data types

In Fig. 6, we present the performances of each of the data forms (*datatypes*) derived from the basic data transformations. CS and QN were removed from further analysis as mentioned previously. Results in the box plots in Fig. 6 show that 4 out of 6 times Log-2 TPM (l2tpm) data type provides best median precision scores. Nevertheless, 2 out of 6 times it provided worst median values. Interestingly, raw data type provided best median score 2 out of 6 times and it only provided worst median value one out of 6 times Considering the largest precision scores that are not outliers, Log-2 TPM provided the best scores 5 out of 6 and Log-2 alone provided it 1 out of 6 times. The pie charts in Fig 6 were plotted for the performance of the data types in only top performing preprocessing combinations. The pie charts in the left column are from C3NET and the ones in the right column from ARACNE using the similar information as in Table 2 and Table 3. The pie charts on the left, for each of the algorithms, show the percentages of the normalization methods from the top 42 ranked preprocessing combinations regarding median rank values. We show more stringent ratios on the right plots by presenting the percentages from top 10 ranked combinations. Considering top 42 median ranks (see Additional file 2), Log-2 TPM appeared in 55%, CPM in 17%, TPM in 14% and Log-2 in 14% of the preprocessing combinations. The second pie chart derived from the top 10 best performing preprocessing combinations of C3NET. Log-2 TPM appeared in 60% in the results. Similar dominant results of Log-2 TPM can be seen in the pie charts of ARACNE in the right column of pie charts in Fig. 6. Regarding box plots and pie charts of Fig. 6, Log-2 TPM appears to be mostly better than others as a basic data type. However, Log-2 and raw data are also useful as they may provide better results too. We do not recommend CPM or Log-2 CPM type data.

**Fig. 6.**
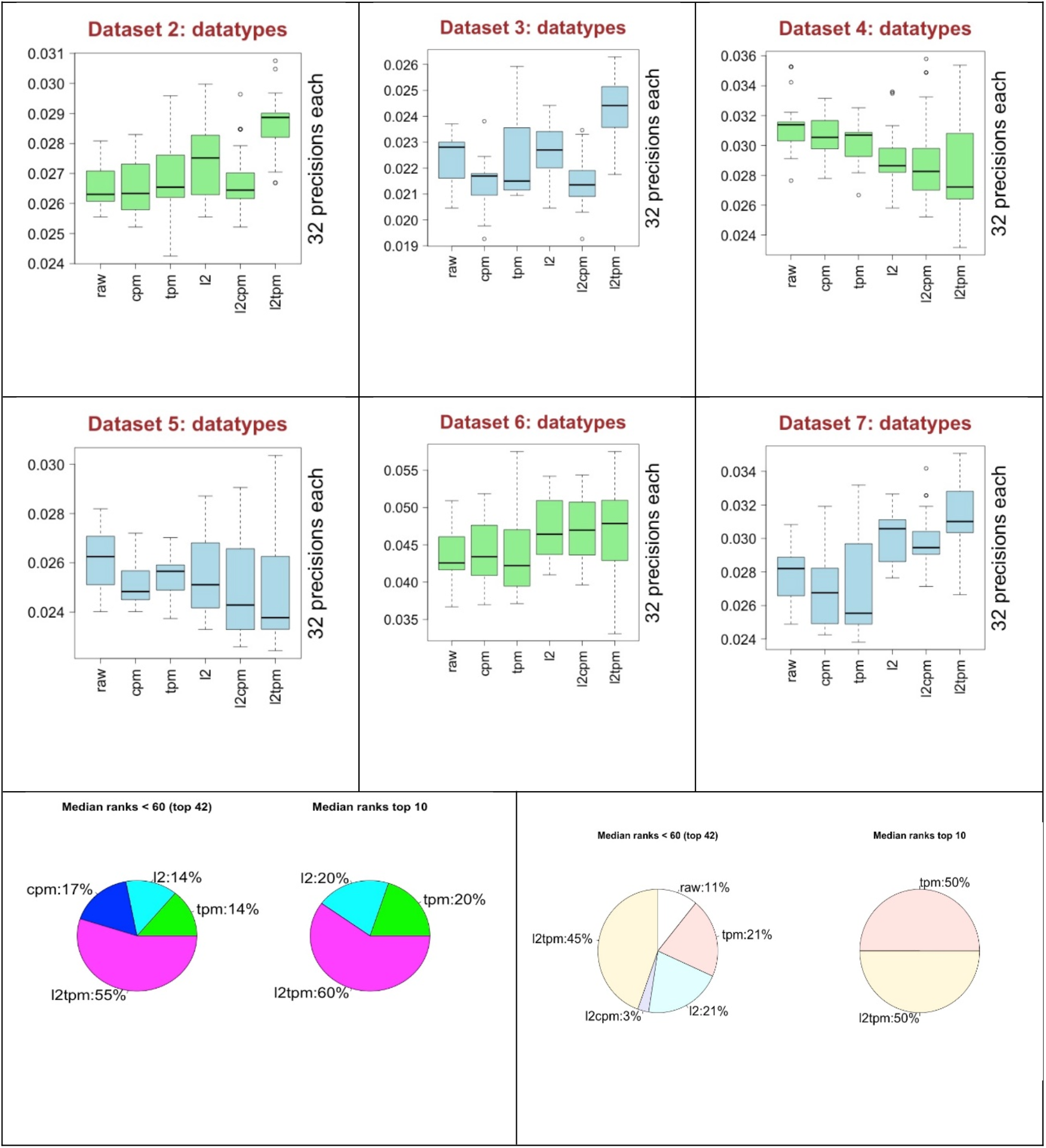
The performance of the data transformation methods (*datatypes*) overall the 6 datasets. Green color is for C3NET and blue color is for ARACNE in the box plots. Pie charts on the left column is from C3NET and the right is from ARACNE.

### Effect of the normalizations methods

In Fig. 7, we see the performance of the normalization methods TMM, VST, RLE and QN along with no normalization applied case (*NoNorm*). Regarding the best performances, there is no one clearly differentiated from others considering all the 6 datasets. Nonetheless, regarding worst performances, QN clearly provides 3 worst median scores and it does not provide a best score over all the 6 datasets. It is a clear evidence of inconsistence performance of QN. This observation lead us to exclude QN from most of the analysis as we have no intention to use it in real applications regarding these outcomes. Considering the median values, VST has 4 highest and 2 lowest ranks. NoNorm has 2 best and 4 worst ranks. It is not safe to choose VST as the best performing normalization method in general as we see it also performs worst in two out of 6 datasets. One thing is worth mentioning that when VST has best median value 4 times, then NoNorm has always the worst values. Additionally, when NoNorm has the best median values twice then VST has the worst median values. This suggest that we need to consider preprocessing combinations with and without VST normalization. RLE and TMM seem to show moderate performances and similar to each other but RLE seem to be slightly better. This analysis was done regarding median performance results but we are normally interested in the best performance scores. As we see from Fig. 7 that the highest performance values, which are not outliers, are showing different ranking than median values. For example, in the result of Dataset 3 in Fig. 7, TMM has the highest score contrary to the rank of its median value. This suggests that we should also consider RLE and TMM for implementations but as they are similar we would suggest to use RLE only.

**Fig. 7.**
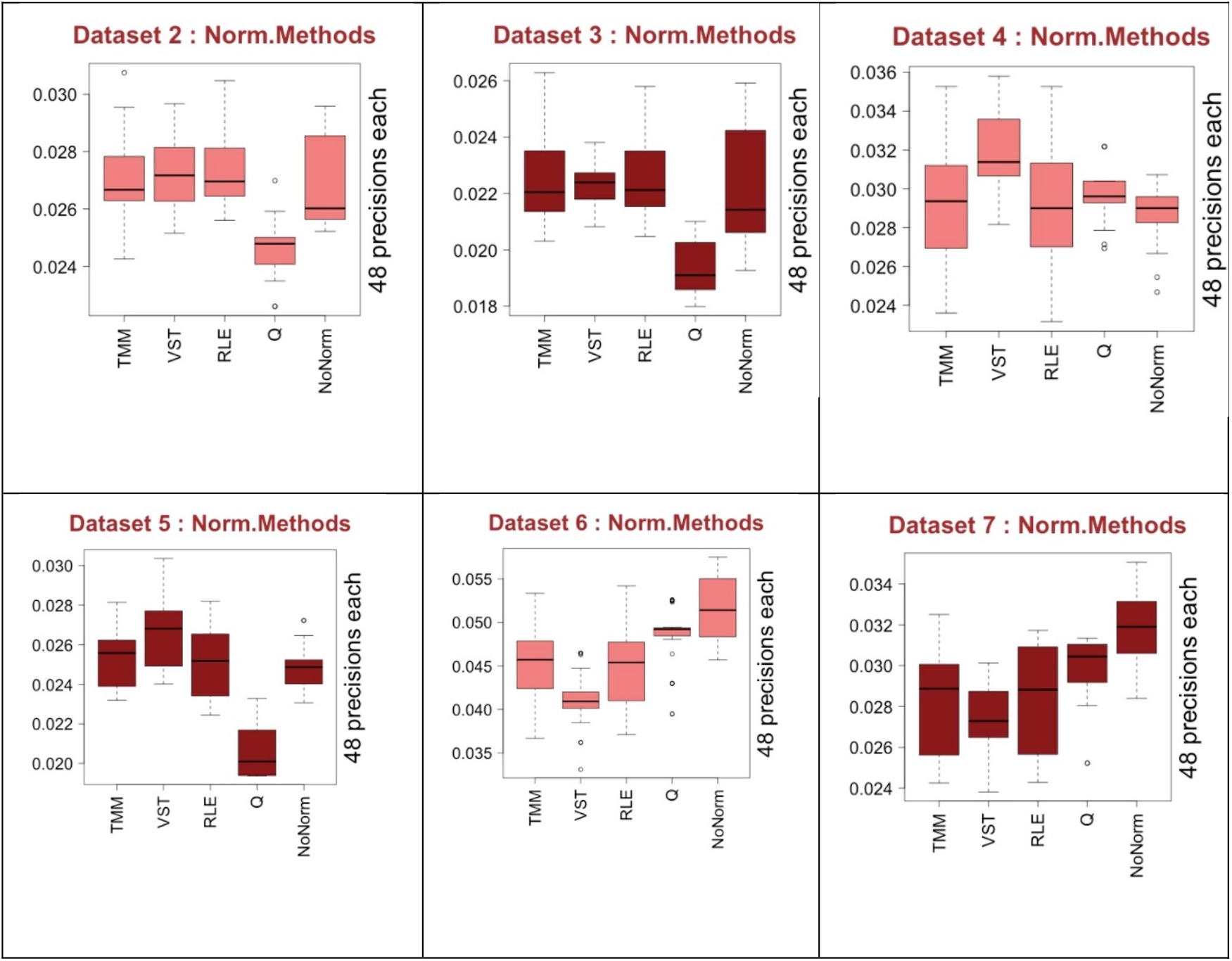
The performance of the normalization methods over all the 6 datasets. Light color is from C3NET and dark color is from ARACNE results.

We have a closer look into the best performing normalization methods in Fig 8. We plot the percentages of the normalization methods in pie charts regarding their median rankings as shown in Table 2 and 3. Upper plots are based on C3NET and lower ones are of ARACNE. Again, the pie charts on the left show the percentages of the normalization methods from the top 42 ranked preprocessing combinations regarding median rank values. The pie charts on right are for top 10 ranked combinations. Regarding the top 42, VST appears for 40%, NoNorm 36%, RLE and TMM 12% in the C3NET results. Regarding the top 10 rankings, NoNorm appears half of the times, then VST for 30%, RLE and TMM for 10%. In the lower plots, we show the results of ARACNE, similarly. In that case, NoNorm dominates more and then TMM, RLE and VST appears with smaller percentages. Considering the top performing combinations, these results suggest primarily not using one of those normalizations. However, VST and RLE should also be tried separately. In general, one of the three options may be tried in preprocessing.

**Fig. 8.**
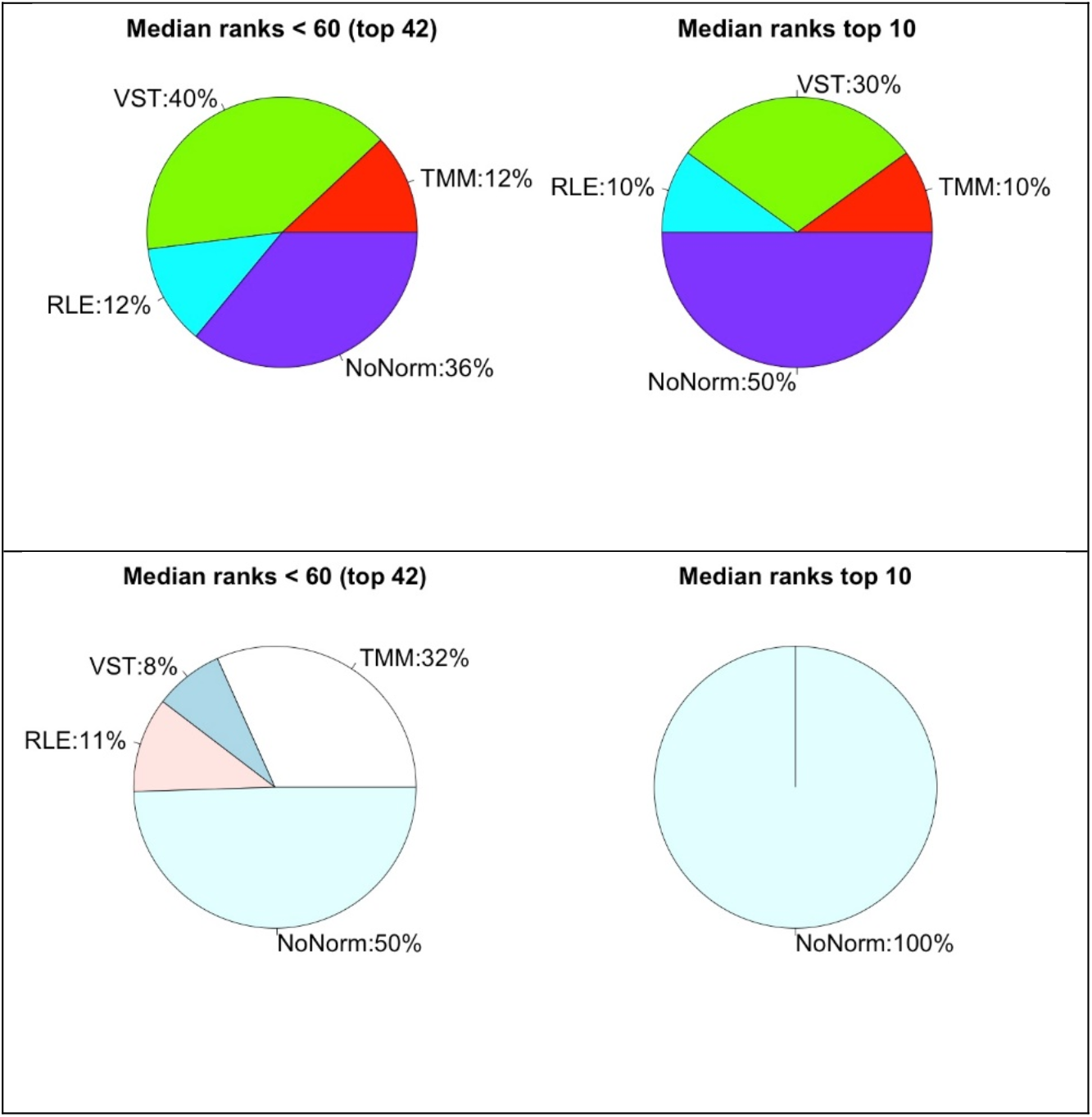
Pie charts that show rank performance ratios of the normalizations methods. Upper part is for C3NET and the lower is for ARACNE results.

### Effect of estimators

In Fig. 9, we present the performances of each of the association estimators. CS and QN were already removed from the analysis. According to the median values, all the estimators provide close performance results. Even for the largest and smallest scores that are not outliers, we do not observe a clear difference mostly among the methods. There are slight differences observed in the performances over all the datasets. This suggest that all the estimators may be selected considering the best performing preprocessing combinations. However, PCC and SCC contain more information as they provide signed correlation values that help to decide the positive and negative associations between genes. B-spline provides mutual information (MI) values that can be only positive values. MI based estimators may capture non-linear relationships too but the performance results do not show a clear empirical evidence in general to favor it.

**Fig. 9.**
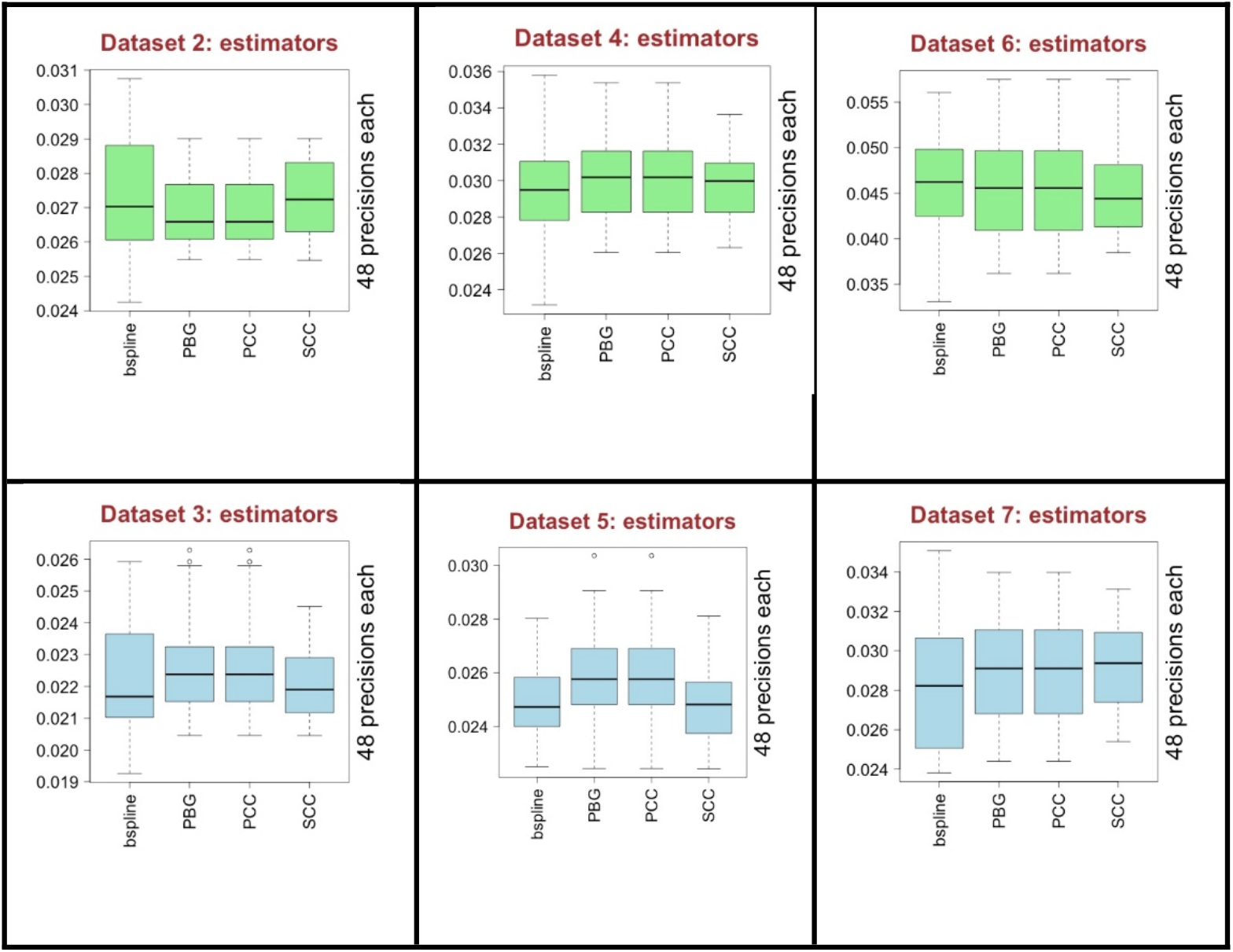
The performance of the association estimators over all the 6 datasets. Green color is for C3NET and blue color is for ARACNE results.

If we look at to the pie charts in Fig 10, presented similar to the previous pie charts, considering the top 42 ranked preprocessing combinations, all of the 4 estimators almost equally performing for both C3NET (upper left) and ARACNE (lower left). Considering the top 10 ranking combinations, B-spline somewhat outperforms the others and PCC and PBG follows it for C3NET (upper right). On the other hand, for ARACNE (lower right), PCC and PBG performs better and B-spline follows it. In general, all the estimators perform somewhat similar as there is no clear difference observed in general.

**Fig. 10.**
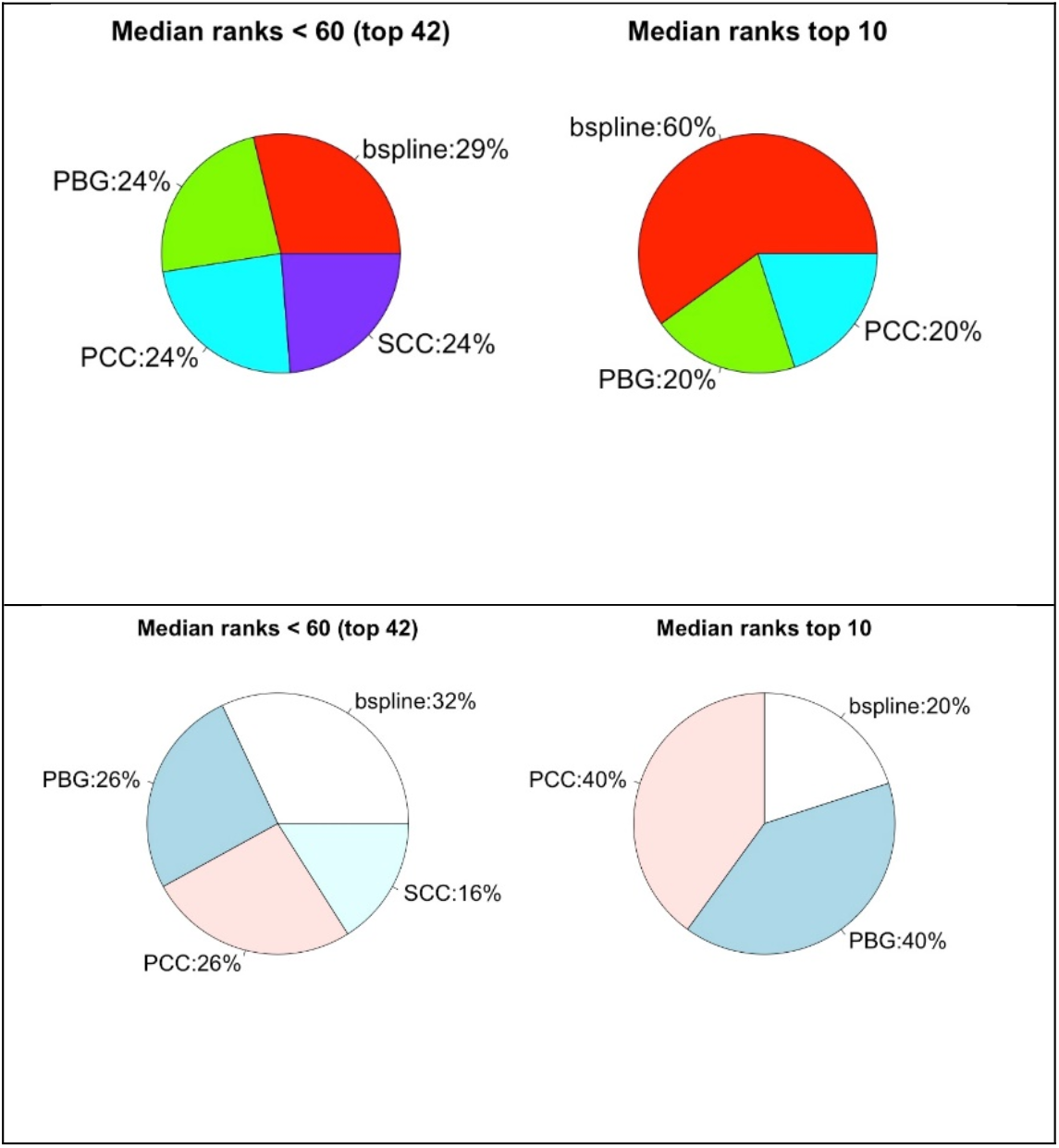
Pie charts that show rank performance ratios of association estimators. Upper part is for C3NET and the lower is for ARACNE results.

### Effect of copula transform (CT)

CT is an important part of GNI over microarray datasets. Here, we investigated its effect on RNA-seq datasets. In Fig 11, the box plots on the left two columns (light color) show the performance of CT in two different datasets using C3NET. Similarly, the other two box plots (dark color) are for ARACNE. Regarding the outcomes of C3NET, not using CT (noCT) seems slightly better. For ARACNE, CT is slightly better regarding the median values but noCT seems better regarding the maximum scores. The pie charts below the box plots are for top ranking preprocessing combinations and provide similar observations. Left column is for C3NET and the right for ARACNE. Although the performance scores of both in general are very close, noCT seems to be giving slightly better scores considering the maximum values and the pie charts. There is no clear difference regarding median values.

**Fig. 11.**
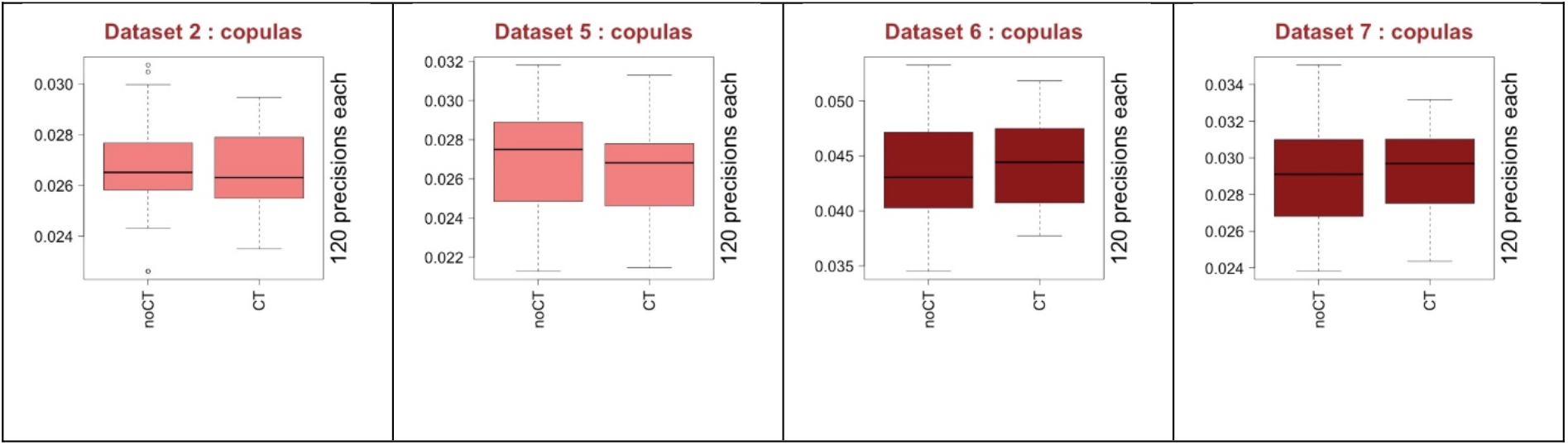
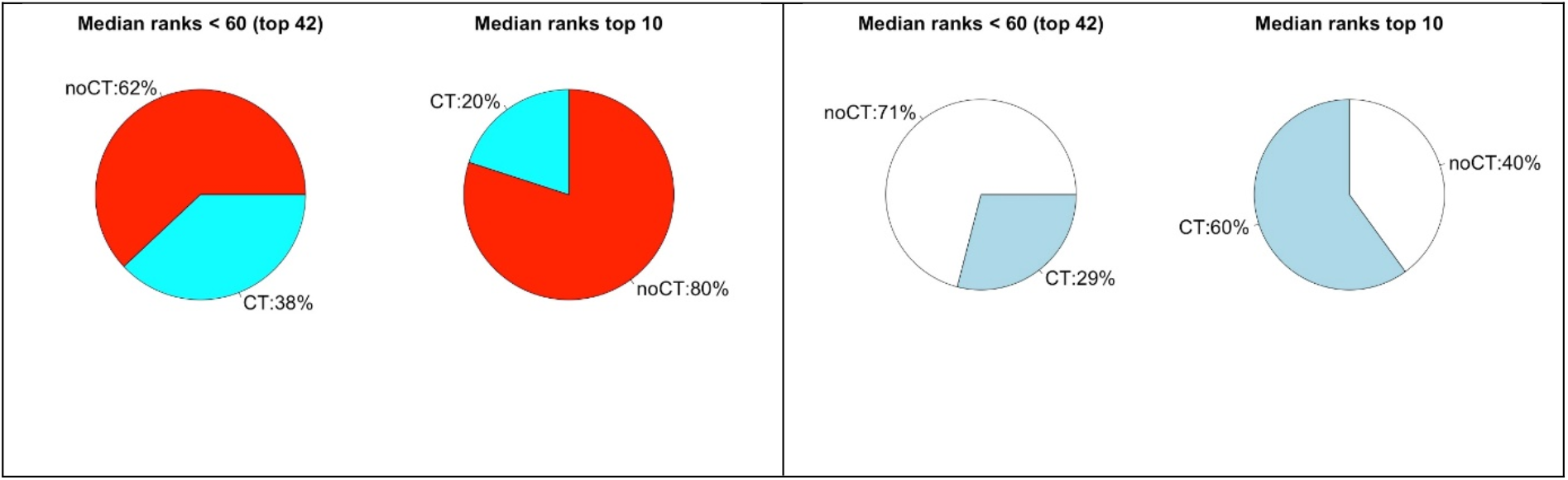
The performance of CT over all the 6 datasets shown by box plots. Left two columns (light color) are from C3NET and the other two (dark color) are from ARACNE results. Pie charts show top ranking performance ratios of CT. Left column is for C3NET and the right is for ARACNE.

### Determining the required sample size for best performances

Our analysis until here focused on the problem of which GNI preprocessing combinations should be chosen and what are the possible consequences if not properly done. Nevertheless, current RNA-seq datasets may be of any size from a few to several hundreds. It is also important to know if the sample size of the available count data is sufficient to attain to the best or optimal GNI performances. This important question was answered in [31] for microarray gene expression datasets and sample size of around 64 was shown as the region where the GNI performances converges dramatically. In this study, we investigated the relation between sample size and GNI performance to determine the required minimum sample size for satisfying performance with RNA-seq datasets. We demonstrate the relations between sample size and performance over various preprocessing combinations and datasets in Fig. 12. In total, we run 288 different simulations for the sample size analysis that helped answering our questions on the issue.

**Fig. 12.**
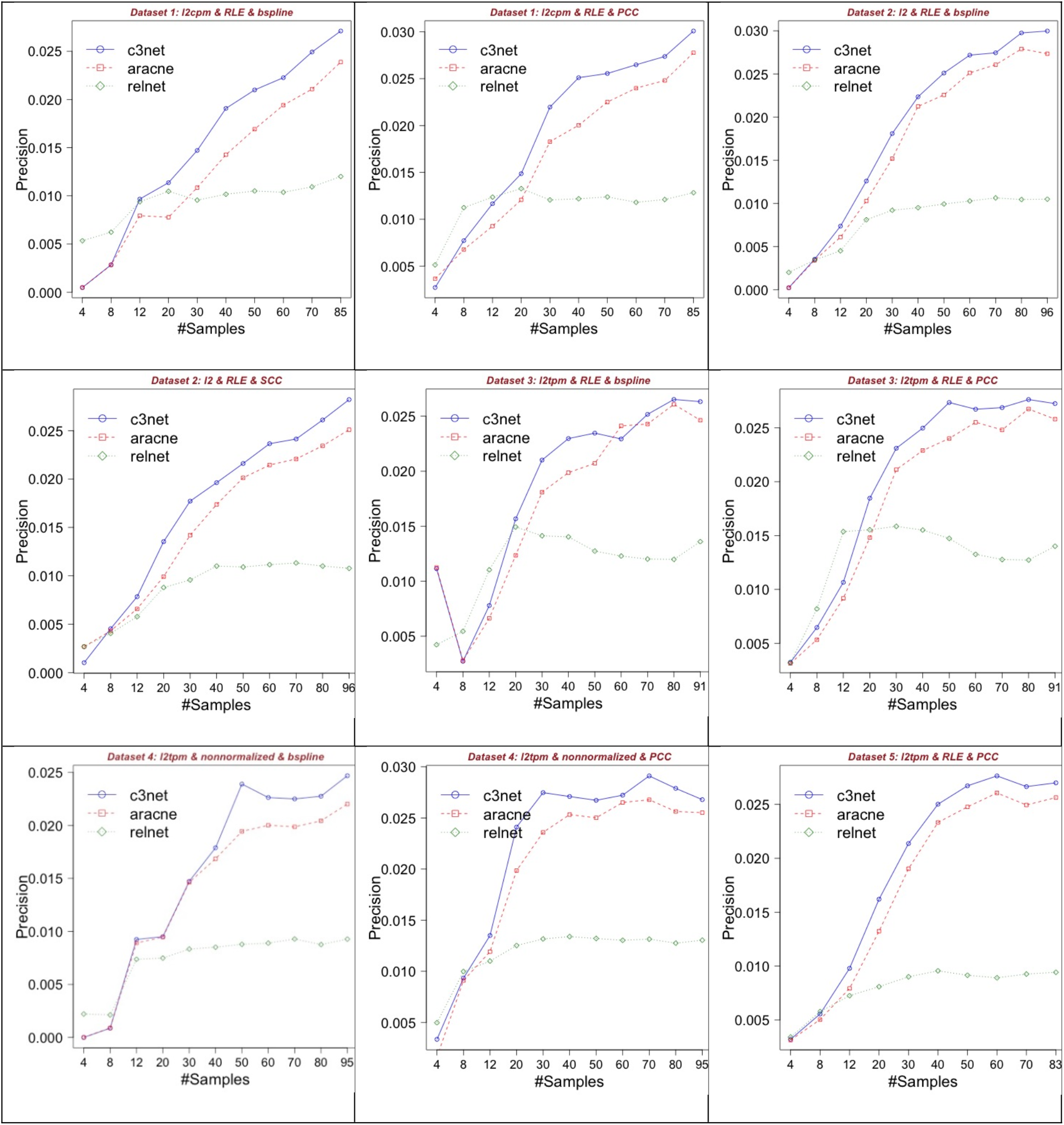
Sample size and performance relationships of GNI preprocessing over RNA-seq data.

Dataset 1 was used as an additional different data set (with sample size 85). It is used for only sample size analysis along with the other datasets. In Dataset 1, using the preprocessing combination B-spline, RLE and Log-2 CPM (first row and column), we observe that there is constant and slow increase in performance until the precision score 0.025 around maximum. C3NET and ARACNE show similar trend until the sample size 85 but C3NET provided the precision score of 0.025 in 70 samples compared to the score for sample size 85 of ARACNE. In the Dataset 1, when we use PCC (first row, second column) along with the same other combination (RLE ad log-2 CPM), then we can obtain the same performance score of around 0.025 in 40 samples with C3NET. With this combination, the performance of C3NET showed somewhat convergence at sample size around 40 but it still produced a performance increase from 0.025 to 0.030 after sample size 70 to 85. ARACNE does not seem to converge but produce a slow performance increase until the sample size 85 and hits the score around 0.027. We do not mention about RELNET (RN) as it was already included the study as a base algorithm and used as a reference to somewhat justify the performance scores of the main representative GNI algorithms ARACNE and C3NET. Because RN is used as the first base step of those and many other algorithms [5].

For Dataset 2, the scores of C3NET from the preprocessing combination B-spline, RLE and Log-2 (first row, third column) start converging to precision 0.027 at sample size 60. The performances show an increase from around 0.027 to 0.03 between the sample sizes 70 to 80. A convergence is observed afterwards until the sample size 96. On the other hand, ARACNE seems to converge at precision around 0.027 after sample size 80. For Dataset 2, using the same preprocessing combination but replacing SCC instead of B-spline (second row, first column), we observe slower increase in performances than before until the sample size 96.

For Dataset 3, the scores of C3NET with the combination B-spline, RLE and Log-2 TPM (second row and column), rapidly converge to precision around 0.022 from sample size 8 to 30. It then increases to the precision score of around 0.026 from sample size 60 to 80 and converges second time afterwards. Here, the scores of ARACNE seem to converge at sample size 60. For Dataset 3, when we replace PCC instead of B-spline in the previous combination (second row, third column), the performance scores of C3NET converge at sample size 50 to the precision of around 0.027. Whereas, ARACNE seem to converge at sample size 60 to the precision of around 0.026.

For Dataset 4, the combination B-spline, no normalization (labeled as *nonnormalized* in these figures) and Log-2 TPM (third row, first column), C3NET and ARACNE converged at sample size 50. On the other hand, using PCC instead of B-spline allowed them to converge at sample size 30 with the similar performance scores. PCC made significant difference with respect to sample size at similar converged performance scores.

For Dataset 5, the combination PCC, RLE and Log-2 TPM provided converged scores at sample size 40 for all the GNI algorithms. Dataset 6 and 7 have smaller sample sizes as 55 and 41, respectively and since their range is small they are not used in this analysis specifically. However, we had compared them in general with all the other datasets considering their full sample sizes in Fig 4 and 5. Dataset 6 and 7 have almost half the sample size of the other datasets. However, they show higher performance scores despite their smaller sample sizes.

Overall, the results suggest that RNA-seq dataset sample size from 30 to 85 may be required to obtain optimal GNI performances. This reflects the inherent heterogeneity of RNA-seq data on the issue of required sample sizes too. Results reveals a very interesting and new conclusion on this topic that we did not see anywhere else before. The critical required sample size number may be as small as around 30 but also as high as around 85. Comparing to microarray datasets, the required sample size was observed to be around 64 for convergence of GNI performance scores [31].

### Filtering effect with variance

Variations in the expression levels are the main components to compute associations scores.

We already filtered RNA-seq data based on only gene expression levels as described in the Methods section. However, we wanted to answer the question whether data filtering based on variation might improve the GNI performance. We filtered out one fifth of the genes that have lowest variance and compared overall performance with the previous results as demonstrated in Fig. 13. In the figure, D label stands for ‘dataset’; numbers refer to the specific Dataset 2 to 7 and HV stands for ‘high variance’ referring to the further filtered datasets with only genes of higher variance. Results show that mostly they get worse if filtered based on variance. Whereas, as seen in Dataset 6 and 7, there may still be potential to get better performance with filtering based on variance. Additional, 3456 analysis (red box plots) were run to make this comparison.

**Fig. 13.**
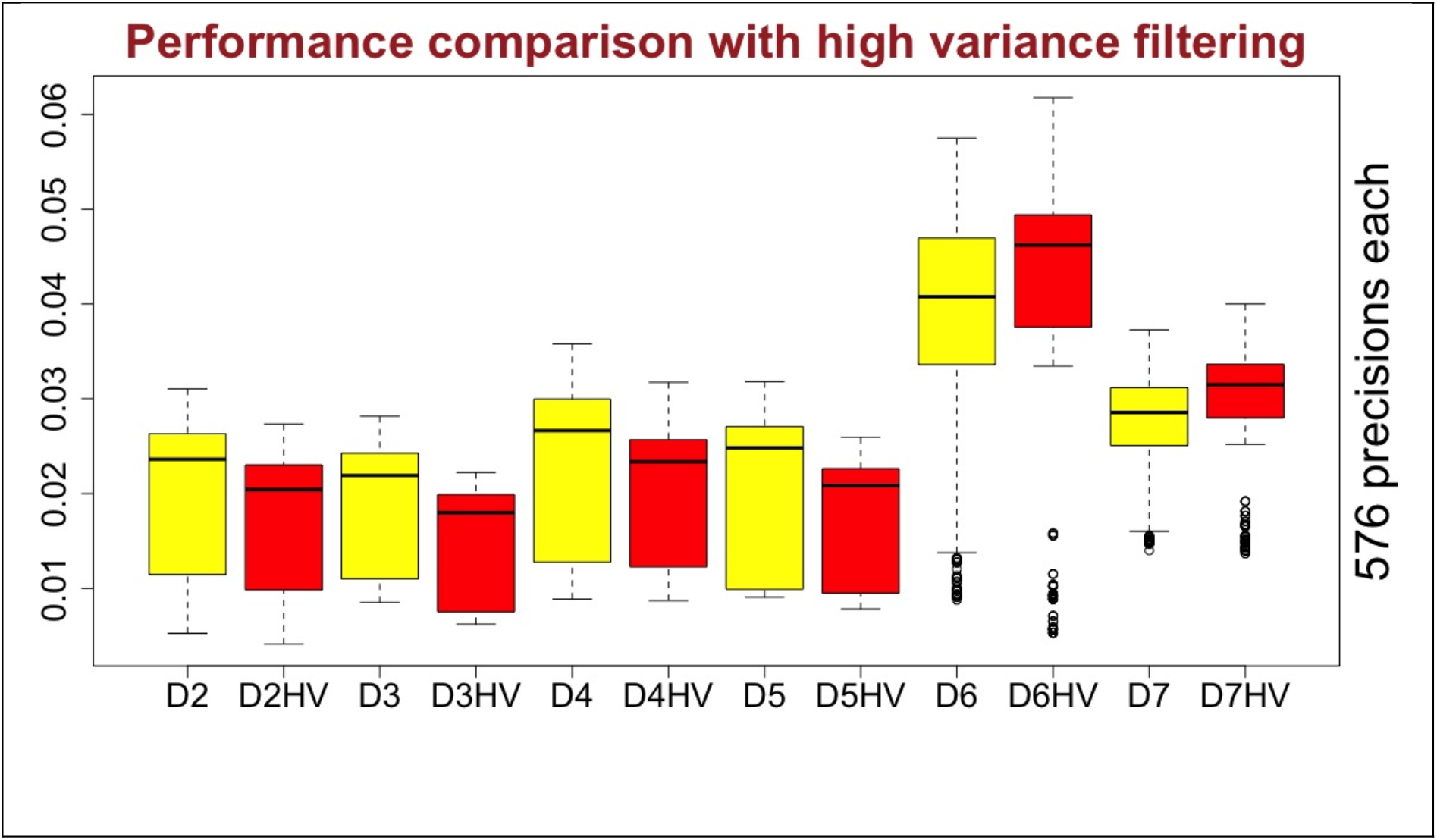
Comparison between default datasets with further filtered versions of them based on variance (HV). D label stands for ‘dataset’; numbers refer to the specific Dataset 2 to 7. HV stands for ‘high variance’ referring to the filtered datasets based on higher variance.

## Discussion

This study helps selecting good preprocessing approaches for the applications of GNI algorithms on RNA-seq datasets. To the best of our knowledge, there is no such systematic study available in the context of GNI over RNA-seq datasets and this study meets an urgent need on the issue. There are very large number of possibilities to combine a preprocessing considering association estimations, normalizations, data transformations (this may also be considered as basic normalizations). Therefore, we selected some popular ones as they are expected to be promising. As illustrated in Fig. 1, initially, the combinations appeared to have 6 different data forms, 5 association estimators, 5 normalizations cases and 2 CT cases. Considering also not implementing some of those methods, in total there were 300 different preprocessing combinations. Considering the three GNI algorithms, we had 900 different combinations that are run over the 6 different RNA-seq datasets, which made 5400 performance analysis to answer the main research question of this study. As we observed very poor performances of CS and QN, we excluded them from further analysis. In the sample size analysis, we run 288 different simulations. In the high variance filtering analysis, we run another 3456 analysis. Total number of analysis presented in this study is 9144. The number of analysis and possible comparisons are overwhelming that limit to include more possible combinations. Other extensive studies may help on this for the similar endeavor in the future. This study also reveals that current preprocessing methods are not satisfactory in the context of GNI over RNA-seq datasets and new approaches are needed that work well in general.

By using 6 different datasets, we found out that there is no best but there are good combinations that work well in mostly but each dataset might need a different preprocessing combination. That directs us to use the literature and perform a quick performance analysis with the good preprocessing combinations and select the best combination specifically for the dataset of interest. Alternatively, one may just choose one of the good ones that we present in this study; knowing the fact that it may not give the best performance but a good one.

Comparing to the performance of randomly generated networks, we observed some close poor performance scores to random networks but we also observed dramatic gains in performance when right preprocessing combinations used. This shows the importance of this kind of studies that will help better selecting the preprocessing combinations to be able to apply GNI algorithms on RNA-seq datasets with some confident. From our extensive analysis, we selected some good preprocessing combinations as specified in the conclusion section.

## Conclusions

We concluded that preprocessing is the most important part of any GNI application over RNA-seq datasets. Because, if not properly chosen, one might end up with very poor results. We also conclude that there is no best preprocessing method but several good combinations for GNI over RNA-seq datasets, as shown in Table 2. We further conclude that the literature interactions databases should be part of any GNI process and be used as a reference to select most suitable preprocessing combination that is specific to the RNA-seq dataset of current interest. For the ease of practical implementation of this, we provided an updated R package, *ganet*, along with this study.

We also concluded that the minimum required sample size may vary from around 30 to 85 for each RNA-seq data for sufficient performances of GNI. We also concluded that GNI algorithm and preprocessing combinations can make significant differences in the required sample size.

For the practical applications in general, instead of selecting a single GNI preprocessing combination, we suggest to select several good ones and apply them for testing their performance with the literature as demonstrated in this study.

Considering all the results, especially Table 1, 2, 3 and Additional file 2, we select the following preprocessing combinations as reference ones among the good ones. First one is the combination of PCC and VST and Log-2 TPM. Second one is PCC and no normalization. The performances of these two can tested and the better one may be selected for faster practical implementation in general. However, in the search of best performance on a specific dataset, then Log-2 TPM or raw should be tried in the combination additionally. If raw data performs better, then Log-2 of the raw should also be tested. Then, VST, RLE and no normalization should be tested in the combinations. Then, PCC, SCC and B-spline should be tested in the combinations. This way 18 possible combinations may be tried in the search of the best performing combinations for a new dataset. This study concluded on two main combinations or 18 other good preprocessing combinations out of 300 possible ones considered in the beginning of this study.

## Methods

### GNI and Preprocessing components

We provided important practical information about the used methods in this section. However, further details are provided for all the methods in Additional file 1. For the GNI as seen at Step 6 in Fig. 1, we have used three GNI algorithm ARACNE [21], C3NET [22] and RELNET [23] with R software packages *minet* [32] and *c3net* [33]. For ARACNE, when we set the DPI parameter to 0.1 as used in their original paper, it infers more than 40000 interactions and performance of it was not good as also seen in [22]. In order to increase its performance, we set DPI parameter to 0. This removed all the weakest edges in the triangles in the network and ARACNE performed dramatically better. However, in this case it predicts very low number of interactions (around 5000) similar to C3NET. If ARACNE will be used with the default DPI parameter 0.1 to be able to infer large numbers of interactions, then it can provide performance results only in between C3NET and RELNET [22]. One must be cautious in using ARACNE with a DPI parameter higher than 0.

At Step 5 in Fig. 1, we used 5 association estimators B-spline of order 2 [26], Chao-Shen [27], Pearson Correlation Coefficient (PCC), Spearman Correlation Coefficient (SCC) and Pearson-based Gaussian (PBG) as described in detail in [24, 28]. Implementations of them were performed with the R software package DepEst that includes functions for 11 different association estimators, which also allows parallel processing [34].

At Step 4 in Fig. 1, For copula transform (CT), we used the function *copula* in c3net software package [33]. At Step 3 in Fig. 1, We used 4 advanced normalizations, Variance-stabilizing transformation (VST) [18] of DeSeq2 R package [19], the trimmed mean of M-values normalization [12] and also relative log expression (RLE) [18] from the *calcNormFactors* function of edgeR [20] R software package and quantile normalization (QN) [11]. Further details of all these methods are provided in the Methods sections of Additional file 1. At Step 2 in Fig. 1, For CPM scaling we followed the procedure of the *cpm* function of the popular edgeR software package [20, 35]. For TPM, we also included gene lengths in this procedure as described in the Additional file 1.

### Practical notes

An important practical note is that, before implementing any of these preprocessing steps, we add one to each value in the dataset matrices, which otherwise may cause erroneous results because of possible zeros in the matrix. Adding one to matrix do not change the results but may prevent from failing because of zeros. Also, if a result is obtained much better than the ones presented in this study, one must be very cautious and check the predicted network. We had come across cases where a few of the genes, who have extremely larger variations than most of the others, may dominate the network as big hubs. This biased case may cause to higher scores if the gene is one of the well-studied ones in the literature. Our default filtering and adding one before each preprocessing helped overcome these cases in the studied datasets. However, one should always check out the network once predicted for such dominated genes and also other unusual structures of the network.

### Filtering

We first filtered out the NA named genes, meaning not annotated genes from the data. If there are multiple rows with the same gene symbol, we kept only one of the raw of that gene with the highest variance. After cleaning up the dataset, we filtered them as follows: We first removed those genes whose CPM expression values less than or equal to 0.1 for at least 20% of the samples. In Fig. 14, upper left figure, we plot CPM expression values of a gene that need to be filtered and this rule eliminates such genes. In Fig. 14, upper right figure, we plot all the genes with respect to this filtering threshold and number of samples. As it is observed in the lower left tail of the plot, there are around 5000 genes that might be filtered out by the specified rule and threshold. CPM value 0.1 corresponds to 2 expressed counts with a lowest sequencing depth (library size) of around 20 million. If the library size is 75 million then it corresponds to approximately 7.5 expressed counts [36]. We further filtered the genes whose maximum CPM value is less than or equal to 0.7 over all the samples. Similarly, we then filtered the genes whose mean CPM value is less than 0.35 over all the samples. These are based on our exploratory analysis as mentioned over Fig. 14. There is no well-established, commonly used and scientifically justified approach on filtering too. This topic needs a separate extensive study in general. The filtered and non-filtered density plots of Dataset 2 is presented in Fig. 14 (second row). This shows that our filtering helped somewhat eliminating poorly expressed genes and the data has a better distribution after filtering.

**Fig. 14.**
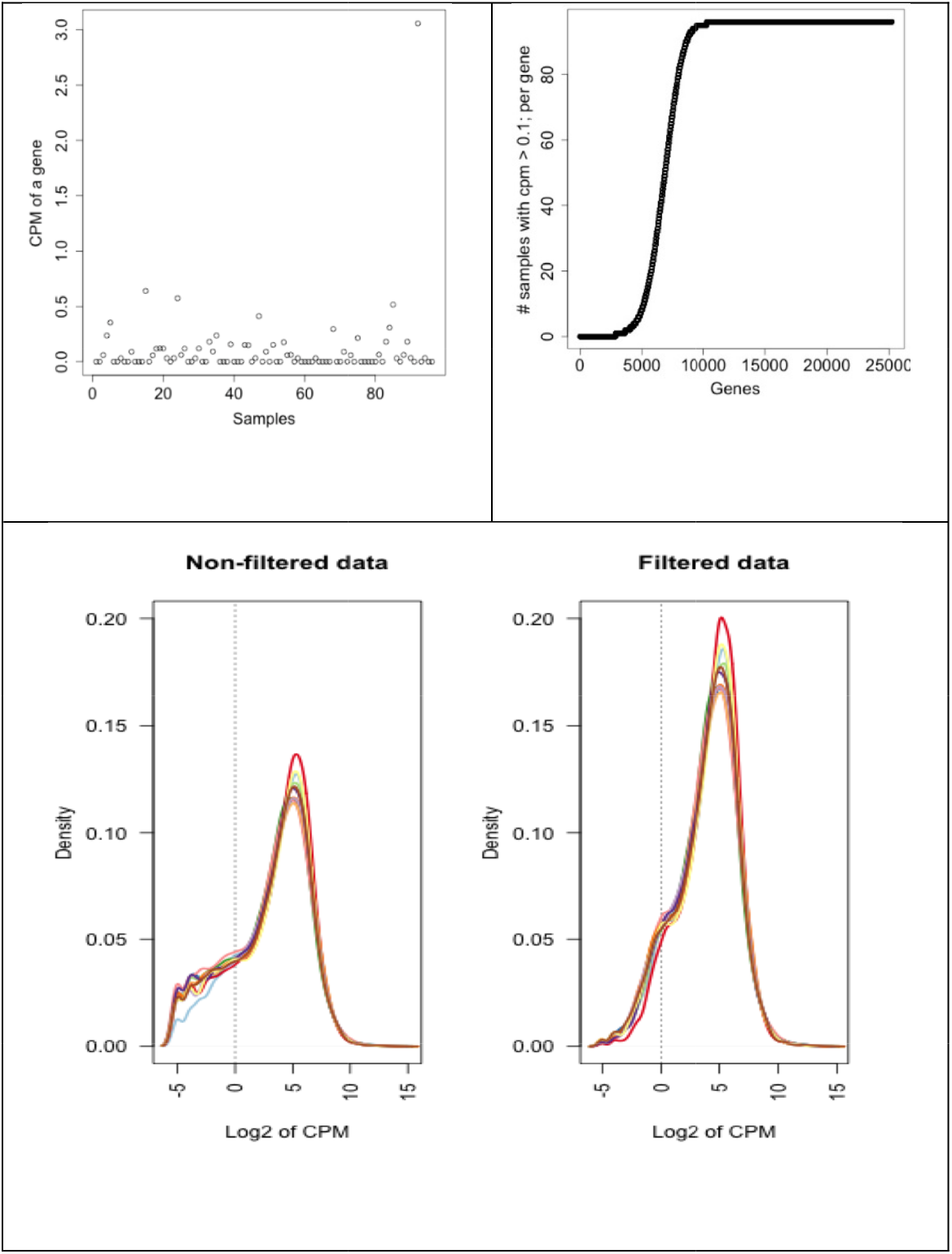
Presented to show the need for filtering. Upper left plot show an example of a gene to be filtered. Upper right shows the number of genes whose CPM < 0.1 relative to number of samples. Below density plot shows both non-filtered and filtered data that is Dataset 2 for all these plots.

### The literature as reference network and performance metrics

In order to assess the performance of GNI results, we used the combination of all the suitable wet-lab validated interaction databases we found in the literature as in [37]. The hypothesis behind the approach is the following: Regardless of cell condition, if there is an experimentally validated gene interaction pair in the literature and if we predict the same gene interaction pair with a statistically significant association score from our datasets then it is very highly likely that the pair can be validated in our cell condition as well. Considering this hypothesis, when a GNI algorithm infers a gene interaction pair and if it is available in the combined database of the literature then we accept that interaction as true positive (TP) in the performance metric [37]. In the other case, when a predicted interaction is not in the literature, it is accepted as false positive (FP). Since the literature is not complete, some of those FP results may in fact be TP. This means, performance is more likely to get better but not worse in the case of an absolute reality. This hypothesis is only accepted based on existence of data; it means positive cases. The predictions are the first positive case, and the interactions in the literature database are second positive case. Since the literature database is far from completeness, when we do not predict an interaction and it is not in the literature database this cannot be accepted as a true result on a negative case (True Negative or TN). Apparently, that interaction may not be in the literature yet because it is not experimentally tested yet in the cell condition of interest. If we use these TN results in the performance metric, then the measured performance might actually be lower with respect to the absolute reality. Therefore, it is not safe to use such cases in the performance metrics with an incomplete reference network such as the literature. Similarly, when we do not predict an interaction and but it is in the literature database this cannot be accepted as a false result on a negative case (False Negative). The main hypothesis to use the literature does not cover those negative cases. Therefore, we mainly measure performances using the precision metric that only consider positive cases (True Positive and False Positive). This may not a perfect metric but when comparing algorithms, all of them face the same biases and we can have reliable conclusions relatively when comparing different algorithms. We explain and discuss more on the performance metrics in detail in Additional file 1.

We tested whether it makes sense to accept the hypothesis of using the literature as a relative reference network for performance evaluation using the randomly generated gene networks and its overlap with the literature. The result was not significant with respect to the overlap analysis. We gave more details on this below while describing random networks. Comparing it with the inferred networks by GNI algorithms, we observed very highly significant overlap results over the literature in general. These tests support the hypothesis to use the literature and we have some level of justification to use the literature as a reference network for globally assessing, or at least comparing, the inference performance of GNI algorithms on real datasets. Nonetheless, we are aware of the fact that when a gene interaction pair is labelled as TP according to the approach, it is not an absolute TP and we might still make mistake. But for the purpose of comparison of various algorithms on real and especially de novo datasets, this idea is very helpful and works well.

The idea of using the literature for GNI performance evaluation is also not new to this study. It was initially used in [37] and since then there are many other studies utilized all or part of the literature for inference or performance assessment of GNI [38–42]. For the practical implementation of it, an R software package ganet (version1.0) was presented in [37] that combined, by 2011, all the available manually curated databases that include experimentally validated interactions. The total number of combined unique interactions was about 550000. The software also provided some functions to run overlap analysis for GNI performance assessment. In this paper, we present a significantly updated version of the R package, ganet (version 2.2) by updating the interactions from existing databases and also by adding more databases as specified below. The literature database in ganet now includes 936850 unique interactions that can be used as a practical reference database for performance assessment. We also added hyper-geometric test function for overlap analysis along with the available Fisher’s exact test function. The updated software package ganet v2.2 is available at a public repository that can be downloaded and used. We utilized from ganet package in our RNA-seq preprocessing performance evaluation pipeline too.

### RNA-seq preprocessing performance evaluation pipeline

Since one of the conclusions of this study is to evaluate the performance each de-novo dataset with the suggested good combinations, we developed a pipeline than can be run efficiently over high performance clusters or on a single computer for a smaller size evaluation. It is based on Snakemake [29], which is an efficient workflow management system. The pipeline allowed evaluating the performance of hundreds of preprocessing combinations approximately in several hours as it runs parallel over clusters. Although it is prepared to evaluate the performance of different preprocessing combinations, if needed, it can also be used as benchmark to evaluate the performance of GNI algorithms as well.

### R software ganet v2.2

The literature database in R software package *ganet (version1.0*) was including the following databases by 2011: BioGRID [43], Human Protein Reference Database (HPRD) [44], Molecular Interaction Database (MINT) [45], IntAct [46] and B-Cell Interactome (BCI) [47]. With this study, we have updated newer version of ganet (version 2.2) from those databases and also added the Database of Interacting Proteins (DIP) [48], innateDB [49], CORUM [50] and the Mammalian Protein-Protein Interaction Database (MIPS) [51] databases, which make 936850 unique interactions in total in the current version. One can access to most of these databases from the server of [52]. We also added two more functions for overlapping analysis, *ganet.hyperg*, which runs hyper-geometric test, and for practical performance assessment, *ganet.getperformance*. The software ganet v.2.2 is available to be downloaded and used from a public repository. The details of how to install and use ganet with a practical example can be seen in Additional file 1.

### Random network performance for reference level

We generated 10 different random networks with 4748 interactions that is the same as the number of predicted interactions of the best scoring preprocessing combination in the best performing dataset, which is Dataset 6. We then performed the same overlap analysis for the random networks as done so far in this study. The literature database in ganet and the random networks overlap analysis resulted average precision of 0.003 and p value of 0.58 from hyper geometric tests. These results are very useful as we can employ them as base reference levels on performances. Any performance result close to or worse than the performance of the random network is considered as very poor and the associated combination is excluded with no doubt. For example, it appeared that the combinations that include CS estimator provided very poor results and excluded from further analysis. In fact, CS estimator was among the good ones in microarray analysis [25]. This shows how important is to choose the right GNI and preprocessing combinations for the analysis of RNA-seq datasets. On the other hand, if right combinations are chosen one can blindly infer meaningful results.

### Datasets

RNA-seq datasets were downloaded conveniently using the *recount* [53] R package and its repository. We provide the details of each dataset here. Each dataset was given a name for the analysis of this study. They are mentioned in this paper with names Dataset 1 to 7 (or Data 1 to 7 in case there is not sufficient space in the figures).

#### Dataset 1

This dataset has GEO accession number GSE79970 [54]. The title of the dataset in GEO is ‘Peripheral Blood Mononuclear Cells (PBMC) Gene Expression-Based Biomarkers in Juvenile Idiopathic Arthritis (JIA)’. After filtering, we had 14398 unique genes and 85 sample in the raw dataset. This dataset was only used for sample size analysis in this study.

#### Dataset 2

The GEO number of this dataset is GSE57148 [55] and its SRA number in recount repository (https://jhubiostatistics.shinyapps.io/recount/) is SRP041538. The title of this dataset in GEO database is ‘Characterizing gene expression in lung tissue of COPD subjects using RNA-seq’. We used 96 of COPD samples of this dataset as Dataset 2. After filtering, there were 17520 unique genes and 96 samples in the raw dataset.

#### Dataset 3

This dataset has the 91 normal samples of GSE57148 from the same publication [55] of Dataset 2. After filtering, there were 17517 unique genes in the raw dataset.

#### Dataset 4

GEO number is GSE54456 [56] and SRA number is SRP035988. The title of this dataset in GEO is ‘Transcriptome analysis of psoriasis in a large case-control sample: RNA-seq provides insights into disease mechanisms’. We used 95 samples that are obtained from lesional psoriatic skin. After filtering, there were 16888 unique genes in the raw dataset.

#### Dataset 5

This dataset has 83 normal samples of GSE54456 from the same publication [56] of Dataset 4. After filtering, there were 17196 unique genes in the raw dataset.

#### Dataset 6

GEO number is GSE69529 and SRA number is SRP059039. The title of this dataset in GEO database is ‘Elucidating the etiology and molecular pathogenicity of infectious diarrhea by high throughput RNA sequencing’. This study is based on whole blood samples. We used 55 of Rotavirus type samples of this dataset for Dataset 6. It is worth reminding that this dataset performed best overall despite its smaller number of samples. After filtering, there were 15040 unique genes in the raw dataset.

#### Dataset 7

This dataset has 41 Salmonella samples of GSE69529 from the same study of Dataset 6. After filtering, there were 15112 unique genes in the raw dataset. Despite its low number of sample, this dataset has second best performance.

## List of abbreviations

BS: B-spline
CPM: counts per million
CS: Chao-Shen
CT: copula transformation
GNI: Gene network inference
GRN: Gene regulatory networks
GSEA: Gene Set Enrichment Analysis ()
Log-2 or l2: Logarithm with base 2
noCT: no copula transformation
NoNorm or none: No normalization (also denoted as *nonnormalized*)
QN or Q: quantile normalization (QN)
PCC: Pearson Correlation Coefficient
SCC: Spearman Correlation Coefficient
PBG: Pearson-based Gaussian
RLE: relative log expression (RLE)
TMM: The trimmed mean of M-values normalization
TPM: transcript per million)
VST: Variance-stabilizing transformation

\# symbol stands for “number of”

## Declarations

The content is solely the responsibility of the authors and does not necessarily represent the official views of the National Institutes of Health.

## Acknowledgements

We used R [17] and Snakemake [29] for the programming. This study was performed in 2019 while Gökmen Altay and Jose Zapardiel-Gonzalo were performing research in La Jolla Institute for Immunology.

## Availability of data and materials

All the data and software used in this study are publicly available as specific in the Methods section.

## Authors’ contributions

GA and BP conceived the study and wrote the paper. GA performed all the analysis, developed and run the software programs. JZ contributed in programming over clusters and some of the analysis. BP coordinated the study and contributed to the analysis. All authors read and approved the final manuscript.

## Consent for publication

Not applicable.

## Ethics approval and consent to participate

Not applicable.

## References

1. Dutta A, Le Magnen C, Mitrofanova A, Ouyang X, Califano A, Abate-Shen C: Identification of an NKX3.1-G9a-UTY transcriptional regulatory network that controls prostate differentiation. Science 2016, 352:1576–1580.

2. Langfelder P, Horvath S: WGCNA: an R package for weighted correlation network analysis. Bmc Bioinformatics 2008, 9.

3. Subramanian A, Tamayo P, Mootha VK, Mukherjee S, Ebert BL, Gillette MA, Paulovich A, Pomeroy SL, Golub TR, Lander ES, Mesirov JP: Gene set enrichment analysis: A knowledge-based approach for interpreting genome-wide expression profiles. Proceedings of the National Academy of Sciences of the United States of America 2005, 102:15545–15550.

4. Altay G, Emmert-Streib F: Revealing differences in gene network inference algorithms on the network level by ensemble methods. Bioinformatics 2010, 26:1738–1744.

5. Emmert-Streib F, Glazko GV, Altay G, de Matos Simoes R: Statistical inference and reverse engineering of gene regulatory networks from observational expression data. Front Genet 2012, 3:8.

6. Nagalakshmi U, Wang Z, Waern K, Shou C, Raha D, Gerstein M, Snyder M: The transcriptional landscape of the yeast genome defined by RNA sequencing. Science 2008, 320:1344–1349.

7. Black MB, Parks BB, Pluta L, Chu TM, Allen BC, Wolfinger RD, Thomas RS: Comparison of microarrays and RNA-seq for gene expression analyses of dose-response experiments. Toxicol Sci 2014, 137:385–403.

8. Lin Y, Golovnina K, Chen ZX, Lee HN, Negron YL, Sultana H, Oliver B, Harbison ST: Comparison of normalization and differential expression analyses using RNA-Seq data from 726 individual Drosophila melanogaster. BMC Genomics 2016, 17:28.

9. Law CW, Chen Y, Shi W, Smyth GK: voom: Precision weights unlock linear model analysis tools for RNA-seq read counts. Genome Biol 2014, 15:R29.

10. Zwiener I, Frisch B, Binder H: Transforming RNA-Seq data to improve the performance of prognostic gene signatures. PLoS One 2014, 9:e85150.

11. Bolstad BM, Irizarry RA, Astrand M, Speed TP: A comparison of normalization methods for high density oligonucleotide array data based on variance and bias. Bioinformatics 2003, 19:185–193.

12. Robinson MD, Oshlack A: A scaling normalization method for differential expression analysis of RNA-seq data. Genome Biol 2010, 11:R25.

13. Li P, Piao Y, Shon HS, Ryu KH: Comparing the normalization methods for the differential analysis of Illumina high-throughput RNA-Seq data. BMC Bioinformatics 2015, 16:347.

14. Dillies MA, Rau A, Aubert J, Hennequet-Antier C, Jeanmougin M, Servant N, Keime C, Marot G, Castel D, Estelle J, et al: A comprehensive evaluation of normalization methods for Illumina high-throughput RNA sequencing data analysis. Brief Bioinform 2013, 14:671–683.

15. Maza E, Frasse P, Senin P, Bouzayen M, Zouine M: Comparison of normalization methods for differential gene expression analysis in RNA-Seq experiments: A matter of relative size of studied transcriptomes. Commun Integr Biol 2013, 6:e25849.

16. Zyprych-Walczak J, Szabelska A, Handschuh L, Gorczak K, Klamecka K, Figlerowicz M, Siatkowski I: The Impact of Normalization Methods on RNA-Seq Data Analysis. Biomed Res Int 2015, 2015:621690.

17. Team RC: R: A language and environment for statistical computing. R Foundation for Statistical Computing. 2016.

18. Anders S, Huber W: Differential expression analysis for sequence count data. Genome Biology 2010, 11.

19. Love MI, Huber W, Anders S: Moderated estimation of fold change and dispersion for RNA-seq data with DESeq2. Genome Biology 2014, 15.

20. Robinson MD, McCarthy DJ, Smyth GK: edgeR: a Bioconductor package for differential expression analysis of digital gene expression data. Bioinformatics 2010, 26:139–140.

21. Margolin AA, Nemenman I, Basso K, Wiggins C, Stolovitzky G, Dalla Favera R, Califano A: ARACNE: An algorithm for the reconstruction of gene regulatory networks in a mammalian cellular context. Bmc Bioinformatics 2006, 7.

22. Altay G, Emmert-Streib F: Inferring the conservative causal core of gene regulatory networks. Bmc Systems Biology 2010, 4.

23. Butte AJ, Tamayo P, Slonim D, Golub TR, Kohane IS: Discovering functional relationships between RNA expression and chemotherapeutic susceptibility using relevance networks. Proceedings of the National Academy of Sciences of the United States of America 2000, 97:12182–12186.

24. Kurt Z, Aydin N, Altay G: Comprehensive review of association estimators for the inference of gene networks. Turkish Journal of Electrical Engineering and Computer Sciences 2016, 24:695–U1401.

25. Kurt Z, Aydin N, Altay G: A comprehensive comparison of association estimators for gene network inference algorithms. Bioinformatics 2014, 30:2142–2149.

26. Daub CO, Steuer R, Selbig J, Kloska S: Estimating mutual information using B-spline functions - an improved similarity measure for analysing gene expression data. Bmc Bioinformatics 2004, 5.

27. Chao A, Shen TJ: Nonparametric estimation of Shannon’s index of diversity when there are unseen species in sample. Environmental and Ecological Statistics 2003, 10:429–443.

28. Olsen C, Meyer PE, Bontempi G: On the impact of entropy estimation on transcriptional regulatory network inference based on mutual information. EURASIP J Bioinform Syst Biol 2009:308959.

29. Koster J, Rahmann S: Snakemake--a scalable bioinformatics workflow engine. Bioinformatics 2012, 28:2520–2522.

30. Collado-Torres L, Nellore A, Kammers K, Ellis SE, Taub MA, Hansen KD, Jaffe AE, Langmead B, Leek JT: Reproducible RNA-seq analysis using recount2. Nature Biotechnology 2017, 35:319–321.

31. Altay G: Empirically determining the sample size for large-scale gene network inference algorithms. let Systems Biology 2012, 6:35–U28.

32. Meyer PE, Lafitte F, Bontempi G: minet: A R/Bioconductor Package for Inferring Large Transcriptional Networks Using Mutual Information. Bmc Bioinformatics 2008, 9.

33. Altay G, Emmert-Streib F: Structural influence of gene networks on their inference: analysis of C3NET. Biology Direct 2011, 6.

34. Altay G, Kurt Z, Altay N, Aydin N: DepEst: an R package of important dependency estimators for gene network inference algorithms. bioRxiv 2017.

35. McCarthy DJ, Chen YS, Smyth GK: Differential expression analysis of multifactor RNA-Seq experiments with respect to biological variation. Nucleic Acids Research 2012, 40:4288–4297.

36. Law CW, Alhamdoosh M, Su S, Smyth GK, Ritchie ME: RNA-seq analysis is easy as 1-2-3 with limma, Glimma and edgeR. F1000Res 2016, 5:1408.

37. Altay G, Altay N, Neal D: Global assessment of network inference algorithms based on available literature of gene/protein interactions. Turkish Journal of Biology 2013, 37:547–555.

38. Olsen C, Fleming K, Prendergast N, Rubio R, Emmert-Streib F, Bontempi G, Haibe-Kains B, Quackenbush J: Inference and validation of predictive gene networks from biomedical literature and gene expression data. Genomics 2014, 103:329–336.

39. Olsen C, Bontempi G, Emmert-Streib F, Quackenbush J, Haibe-Kains B: Relevance of different prior knowledge sources for inferring gene interaction networks. Frontiers in Genetics 2014, 5.

40. Han H, Shim H, Shin D, Shim JE, Ko Y, Shin J, Kim H, Cho A, Lee EKT, Kim H, et al: TRRUST: a reference database of human transcriptional regulatory interactions. Scientific Reports 2015, 5.

41. Chen GC, Cairelli MJ, Kilicoglu H, Shin D, Rindflesch TC: Augmenting Microarray Data with Literature-Based Knowledge to Enhance Gene Regulatory Network Inference. Plos Computational Biology 2014, 10.

42. Linde J, Schulze S, Henkel SG, Guthke R: Data-and Knowledge-Based Modeling of Gene Regulatory Networks: An Update. Excli Journal 2015, 14:346–378.

43. Stark C, Breitkreutz BJ, Reguly T, Boucher L, Breitkreutz A, Tyers M: BioGRID: a general repository for interaction datasets. Nucleic Acids Research 2006, 34:D535–D539.

44. Prasad TSK, Goel R, Kandasamy K, Keerthikumar S, Kumar S, Mathivanan S, Telikicherla D, Raju R, Shafreen B, Venugopal A, et al: Human Protein Reference Database-2009 update. Nucleic Acids Research 2009, 37:D767–D772.

45. Licata L, Briganti L, Peluso D, Perfetto L, Iannuccelli M, Galeota E, Sacco F, Palma A, Nardozza AP, Santonico E, et al: MINT, the molecular interaction database: 2012 update. Nucleic Acids Research 2012, 40:D857–D861.

46. Kerrien S, Aranda B, Breuza L, Bridge A, Broackes-Carter F, Chen C, Duesbury M, Dumousseau M, Feuermann M, Hinz U, et al: The IntAct molecular interaction database in 2012. Nucleic Acids Research 2012, 40:D841–D846.

47. Wang K, Nemenman I, Banerjee N, Margolin AA, Califano A: Genome-wide discovery of modulators of transcriptional interactions in human B lymphocytes. Research in Computational Molecular Biology, Proceedings 2006, 3909:348–362.

48. Salwinski L, Miller CS, Smith AJ, Pettit FK, Bowie JU, Eisenberg D: The Database of Interacting Proteins: 2004 update. Nucleic Acids Research 2004, 32:D449–D451.

49. Breuer K, Foroushani AK, Laird MR, Chen C, Sribnaia A, Lo R, Winsor GL, Hancock REW, Brinkman FSL, Lynn DJ: InnateDB: systems biology of innate immunity and beyond-recent updates and continuing curation. Nucleic Acids Research 2013, 41:D1228–D1233.

50. Ruepp A, Waegele B, Lechner M, Brauner B, Dunger-Kaltenbach I, Fobo G, Frishman G, Montrone C, Mewes HW: CORUM: the comprehensive resource of mammalian protein complexes-2009. Nucleic Acids Research 2010, 38:D497–D501.

51. Pagel P, Kovac S, Oesterheld M, Brauner B, Dunger-Kaltenbach I, Frishman G, Montrone C, Mark P, Stumpflen V, Mewes HW, et al: The MIPS mammalian proteinprotein interaction database. Bioinformatics 2005, 21:832–834.

52. Cerami EG, Gross BE, Demir E, Rodchenkov I, Babur O, Anwar N, Schultz N, Bader GD, Sander C: Pathway Commons, a web resource for biological pathway data. Nucleic Acids Research 2011, 39:D685–D690.

53. Collado-Torres L NA, Kammers K, Ellis SE, Taub MA, Hansen KD, Jaffe AE, Langmead B, Leek JT.: Reproducible RNA-seq analysis using recount2. Nature Biotechnology 2017.

54. Wong LP, Jiang KY, Chen YM, Hennon T, Holmes L, Wallace CA, Jarvis JN: Limits of Peripheral Blood Mononuclear Cells for Gene Expression-Based Biomarkers in Juvenile Idiopathic Arthritis. Scientific Reports 2016, 6.

55. Kim WJ, Lim JH, Lee JS, Lee SD, Kim JH, Oh YM: Comprehensive Analysis of Transcriptome Sequencing Data in the Lung Tissues of COPD Subjects. International Journal of Genomics 2015.

56. Li BS, Tsoi LC, Swindell WR, Gudjonsson JE, Tejasvi T, Johnston A, Ding J, Stuart PE, Xing XY, Kochkodan JJ, et al: Transcriptome Analysis of Psoriasis in a Large Case-Control Sample: RNA-Seq Provides Insights into Disease Mechanisms. Journal of Investigative Dermatology 2014, 134:1828–1838.

