## Additional file 1 for "RNA-seq preprocessing and sample size considerations for gene network inference"

### Determining optimal data preprocessing approaches and sample size requirements for gene network inference using RNA-seq data

La Jolla instiue for Immunology

*9420 Athena Circle, La Jolla, CA 92037, USA*

G.Altay, J. Zapardiel, B. Peters

Contact:

#### METHODS

We summarize the methods used in this study and refer the readers to the given references in the main text for further details.

##### Gene Network Inference Algorithms:

**RELNET**: Relevance Networks is the first and base algorithm of many information theory based algorithms such as ARACNE and C3NET. It simply calculates association matrix and eliminates the statistically non-significant scores by setting a p-value on the null distribution. We have only kept the most 0.5% of all the association scores after this step for all the analysis. As a typical example, if there are 15000 unique genes in a dataset, this corresponds to 562462 significant association scores out of all the 112492500 association scores. This approach helped the inferred GRNs comparable over all the analysis.

**ARACNE**: uses RELNET in the first step and then in the second step, from the association matrix of significant interactions, weakest edges of all triplets (triangle shaped interactions) are removed if they are below a certain threshold based on a parameter called DPI. In the original paper of ARACNE, they set it as 0.1 but when we set that number then ARACNE was not performing as good as C3NET, therefore we set it to 0 and removed all the weakest edges of the triplets in our analysis. In this case ARACNE resulted small number of predicted interactions similar to C3NET. If more interactions wanted to predict with ARACNE, the DPI parameter may be increased knowing the fact that performance decreases.

**C3NET**: uses RLENET as the first step the in the second step it inly infers the maximum association scored gene pair for each gene. C3NET aims to infer the conservative causal core of gene networks for the sake of highest accuracy instead of inferring the whole network.

**Association estimators:**

**Pearson correlation coefficient (PCC)** calculates linear relationship between two random variables, which therefore can be classified as a parametric or linear and correlation estimator. It does not capture nonlinear relationships.

**Pearson Based Gaussian (PBG),** also called Parametric Gaussian Estimator, estimates mutual information values based on Pearson's correlation with the assumption that the joint distribution is normal. In theory, it can capture linear and nonlinear relationships as a mutual information value.

**Spearman Correlation Coefficient (SCC)** is similar to PCC but the data is ranked before calculating the coefficient.

PCC, PBG and SCC showed very similar GNI performances, which is not surprising considering their similarity.

**B-spline** is based on spline functions and a nonparametric estimator as there are not any assumptions about the data. The spline order $k$ determines the number of bins each of the data points is assigned to. For example, a spline order $k = 3$ means each data point is assigned to three bins in probability calculations of the mutual information value.

**Chao-Shen estimator** combines two different approaches, Horvitz-Thompson estimator and Good-Turing correction of ML estimator.

**Copula Transformation (CT)** converts the data based on ranked values about the range between 0 and 1.

**Counts Per Million (CPM)** or Transcripts Per Million (TPM) can be easily implemented by the *cpm* function of the popular edgeR software package. Basically, it divides the count of each sample to the sum of counts of each sample and multiplies by a million. This in theory prevent the effect of the variations in the library sizes of the samples.

**Normalizations**:

**The trimmed mean of M-values (TMM)** normalization is used as default in the edgR R package that is popular for DE analysis of RNA-seq datasets. It computes a normalization factor for each gene that is applied to library size [1].

**Relative Log Expression (RLE)** implemented using edgeR package and it is the equivalent of the normalization in the DESeq R package that is also popular in RNA-seq DE analysis [1].

**Variance Stabilizing Transformation (VST)** implemented with the popular DESeq2 R package. It transforms data on the log2 scale, which has been normalized with library size. VST generates approximately homoscedastic data that means having constant variance independent of varying mean values. Since VST need integer inputs, before inputting the values to VST function, we multiply the values by 10000 and round them when we take the Log2 of raw scale values.

**Quantile normalization (QN)** is mostly used in microarray gene expression datasets.

QN aims to make the distribution of gene expressions for each array in a set of arrays the same. We implemented QN with preprocessCore R package [2].

**Statistical metrics to evaluate the performance of GNI methods**

The most widely employed statistical measures for the performance assessment of GNI algorithms are precision, recall (sensitivity), specificity, accuracy and F-score [3]. They are all derived from the following ratios: True positive (TP), when the prediction is in the reference network, false positive (FP), when the prediction is not in the reference network, true negative (TN) when a non-predicted edge is not in the reference network and finally false negative (FN) when a non-predicted edge is in the reference network. Their formulas are as follows: precision = P = TP / (TP+FP), recall = R = TP / (TP+FN), specificity = TN / (TN+FP), accuracy = (TP+TN) / (TP+TN+FP+FN), F-score = 2xPxR / (P+R).

According to the hypothesis of using the literature, we accept the predictions in the literature as TP and if not in the literature as FP. We do not make any other assumptions and do not measure anything about the non-predicted edges. Because we know that the literature is incomplete and contains the interactions of all the cell conditions. Therefore, any metric that contains negative ratio is not suitable to asses based on the literature. We are also aware that TP and FP results are not the absolute correct measurements but they relatively give an estimate on the performance. The only metric that does not include negative ratio is precision and used as the main metric for performance evaluation of the analysis. Precision tells us how accurate is the result based on only the predicted set of interactions. It does not give an absolute accuracy measurement but provide a relative one, which appears to be the most suitable metric to use with the literature information. Along with the precision, we also measured the p-value based on hyper-geometric test that measures statistically how significant is the number of overlap with the literature. However, as demonstrated from the results, using this measure alone may be misleading. Because it favors the GNI algorithm that predicts largest number of interactions, which is RELNET (RN) that is already well known to provide worst GNI performance, which was also confirmed in this study. This may be because the formula to get the p-value is biased to the number of TP. RN predicts about 400000 interactions and not surprisingly gives the highest number of TP along with the highest and huge amount of FP. Although RN mostly gets lowest p-value it also has the lowest precision value. Consequently, we used precision as the main performance assessment metric and p-value as the supporting metric to see whether the number of TP predictions of an algorithm is statistically significant or not.

Because whatever the cell condition is, we compare the result from it with the whole literature, which is the combination of almost all cell conditions, and on average all the results are expected to make similar amount of mistake on average with respect to the literature. This approach allows to make comparison of any GNI algorithm on any real dataset, which is not possible otherwise. Most of the studies uses pathway analysis for the evaluation of the inferred networks but it is apparent that it can only be complementary to our approach. Because pathway analysis shows how relevant the gene sets, but not gene interaction pairs, to all the pathways. The relevance of the resulting pathways to the current cell condition of the dataset accepted as validation of the results. Pathway analysis approach is best suited for the results of DE analysis but not GNI. However, considering that GNI results assessed with the literature as we show in this study, pathway analysis may be used as complementary analysis to observe the relevance of the genes in the network to the current cell condition. As this topic is not the focus of this study we end the argument here and leave it for other studies to investigate on it in more detail.

**ganet: R software package for practical access and usage of the literature**

ganet is currently available to download at github. You can easily install ganet from within R session as follows:

```
install.packages("https://github.com/altayg/ganet/raw/master/ganet_2.2.tar.gz",
                        type="source", repos=NULL)   #to install once

library(ganet)   #to load to R session to use
```

ganet v.2.2 incorporates 936850 unique interactions that has been manually curated and experimental support by the literature, which is obtained by combining BioGrid, CORUM, DIP, HPRD, innateDB, IntAct, MINT and MPI databases. Their references can be seen in the main paper. If you want to use the whole combined database by running alldatabases <- ganet.combine() in R, you can have them all in one object; then you also need to cite all these databases along with this paper. But, there is also option to select some of them in the ganet.combine function by running for instance:

someofdatabses <- ganet.combine(BIOGRID = 1, CORUM = 0, DIP = 0, HPRD = 1, INNATEDB = 0, INTACT = 0, MINT = 0, MPI = 0)

Then only BioGrid and HPRD are combined and citation of those are sufficient along with this paper.

In order to detect the overlapping of the predicted network with the literature the you can run the following: overlaps <- ganet.ComLinks(predictedNetwork, alldatabases). Make sure that the first predicted network matrix has at least two columns and its first and second columns have gene pairs of each interaction. For further usage of ganet for statistical overlapping analysis using hyper-geometric or Fisher's exact test, the package help files have detailed information. Below, we present a very simple usage of the ganet package for performance evaluation.

**Tutorial to run ganet R package:**

```
# Following code can be tried with a simple copy paste action in an R session.

# install ganet directly from web within R session
install.packages("https://github.com/altayg/ganet/raw/master/ganet_2.2.tar.gz",
                        type="source", repos=NULL)

#to load to the current R session to use
library(ganet)

# you can use your own predicted network as to column matrix or for trial run the following
# for the example of the predicted network.

data(ganet.ex.net)

# load the unique genes list as a vector for the example predicted network. In the case of a
# real application, this gene list must correspond to the genes when initial analysis started
# from the row dataset.

data(ganet.ex.genes)


#this returns a vector that contains all the performance metrics such as precision and p-val.

res <- ganet.getperformance(PredictedNet=ganet.ex.net, allgenes=ganet.ex.genes)
```

Fig. S1. Some of the interesting RNA-seq data distributions in various forms of Dataset 2.

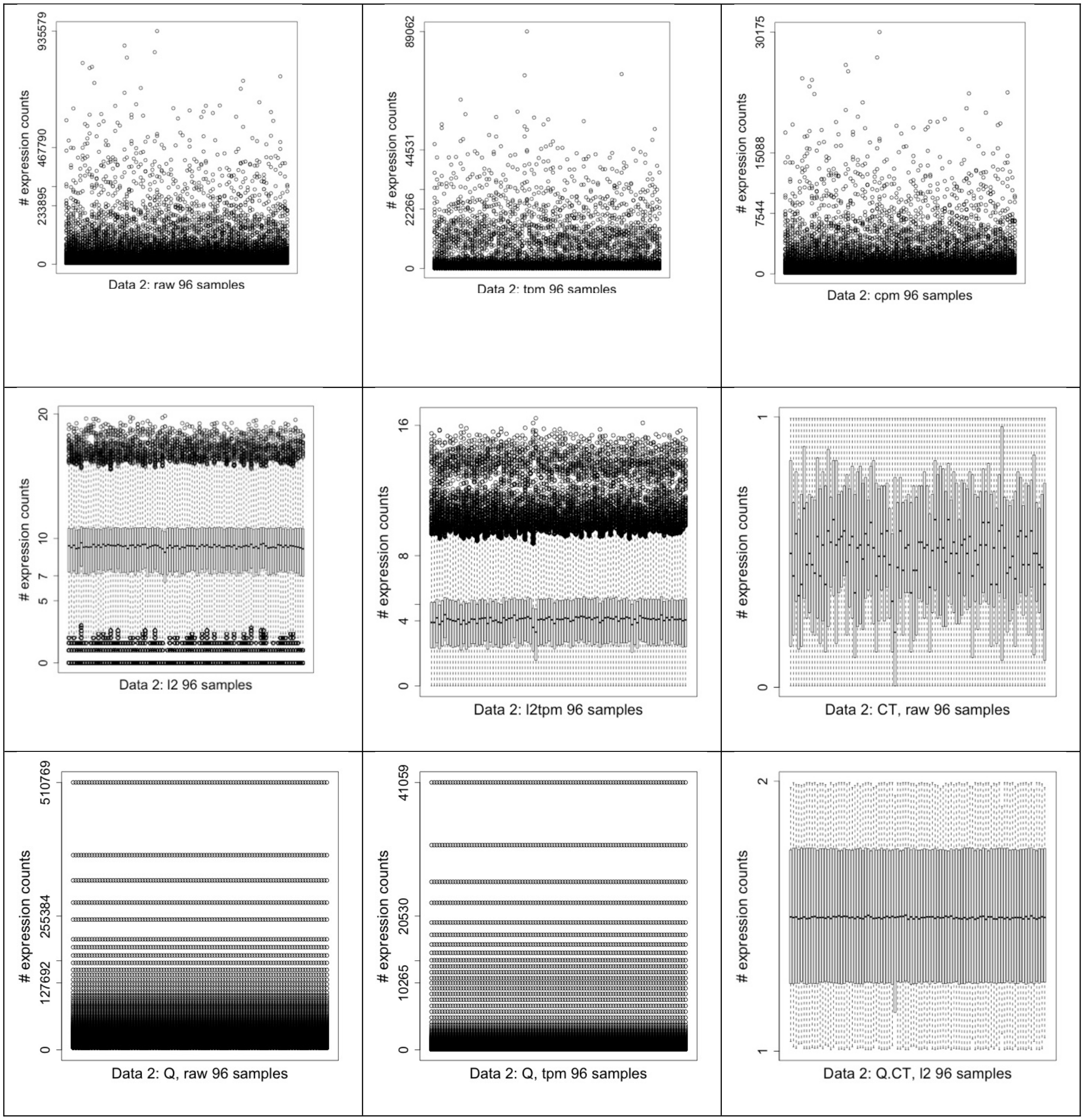

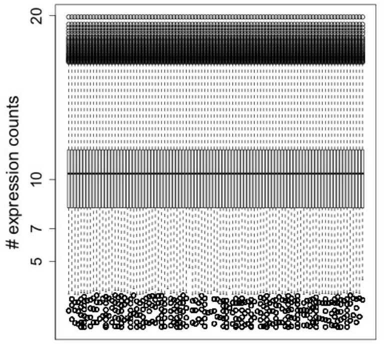

Data 2: Q, l2 96 samples

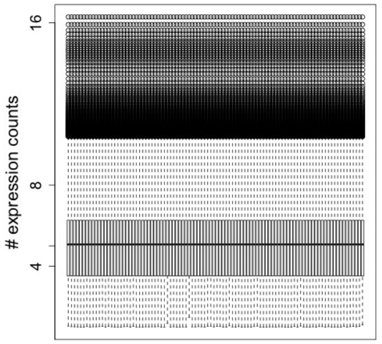

Data 2: Q, l2tpm 96 samples

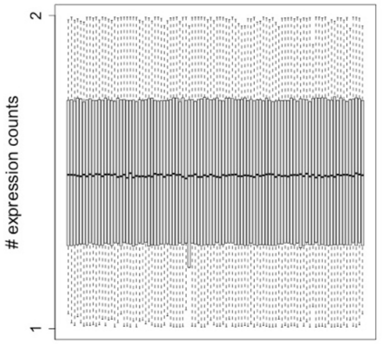

Data 2: Q.CT, l2tpm 96 samples

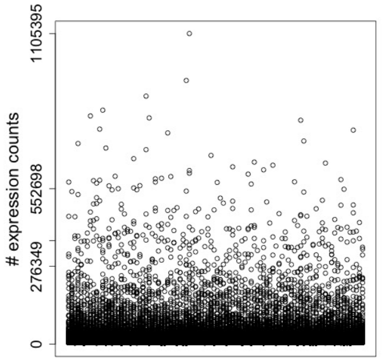

Data 2: RLE, raw 96 samples

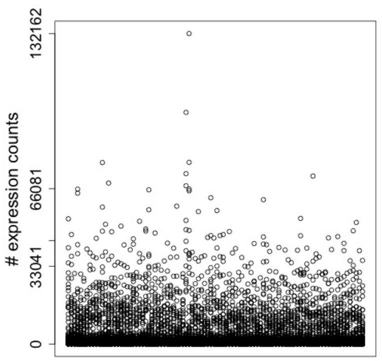

Data 2: RLE, tpm 96 samples

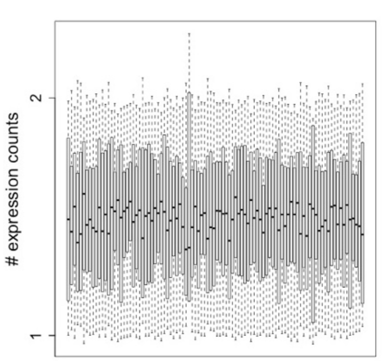

Data 2: RLE.CT, l2 96 samples

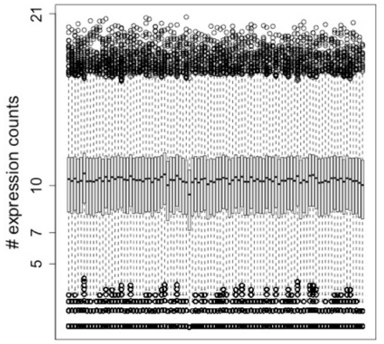

Data 2: RLE, l2 96 samples

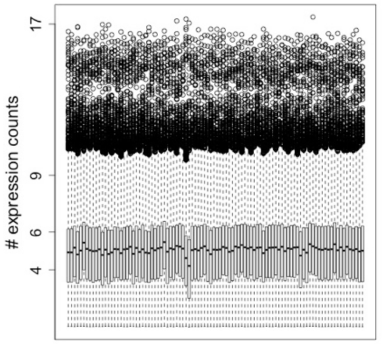

Data 2: RLE, l2tpm 96 samples

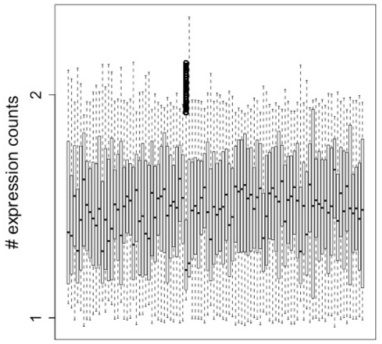

Data 2: RLE.CT, l2tpm 96 samples

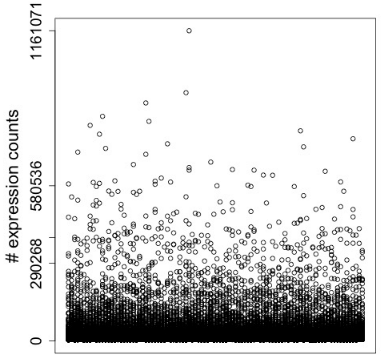

Data 2: TMM, raw 96 samples

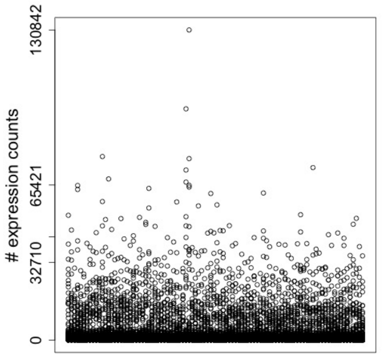

Data 2: TMM, tpm 96 samples

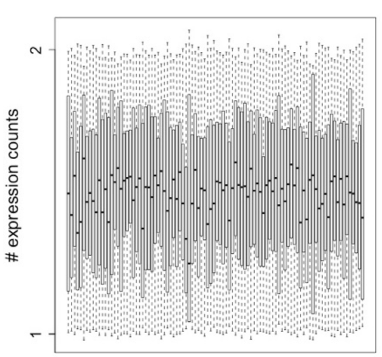

Data 2: TMM.CT, l2 96 samples

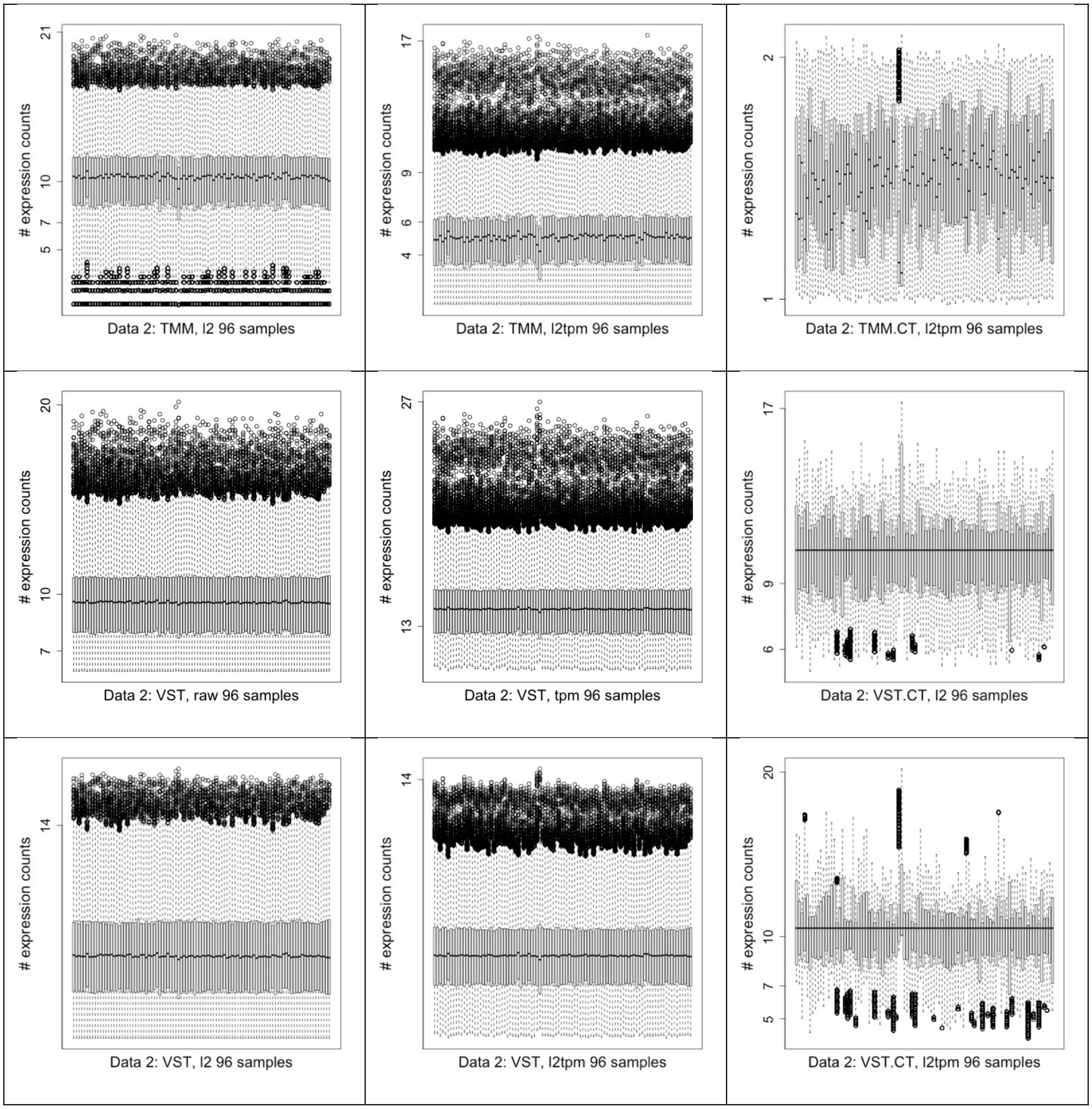
21
10
7
5
### expression counts
Data 2: TMM, l2 96 samples
17
9
6
4
### expression counts
Data 2: TMM, l2tpm 96 samples
2
1
### expression counts
Data 2: TMM.CT, l2tpm 96 samples
20
10
7
### expression counts
Data 2: VST, raw 96 samples
27
13
### expression counts
Data 2: VST, tpm 96 samples
17
9
6
### expression counts
Data 2: VST.CT, l2 96 samples
14
### expression counts
Data 2: VST, l2 96 samples
14
### expression counts
Data 2: VST, l2tpm 96 samples
20
10
7
5
### expression counts
Data 2: VST.CT, l2tpm 96 samples

Fig. S2. Precision performances of all the datasets for all the combinations as horizontal bar plots.

Dataset 3

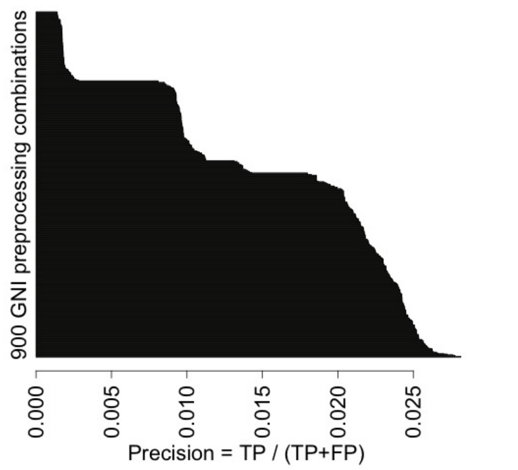


Dataset 4

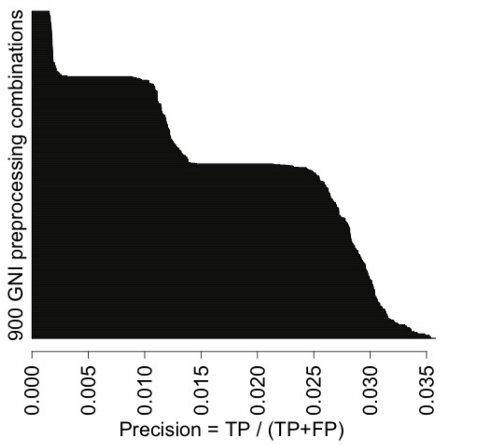


Dataset 5

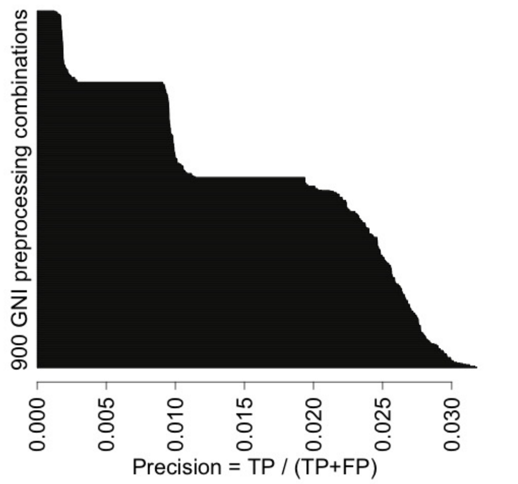


Dataset 7

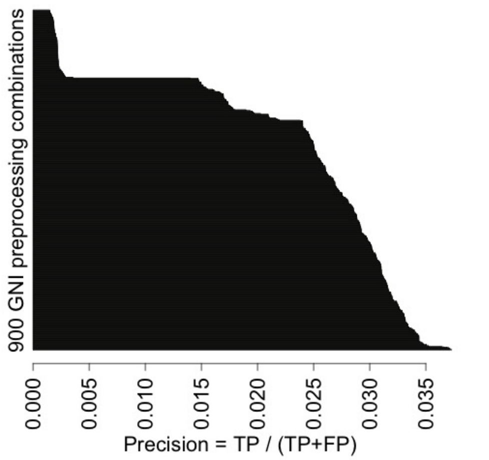

**References for Additional file 1:**

[1] Reddy, R.:A **Comparison of Methods: Normalizing High-Throughput RNA Sequencing Data**. 2015, bioRxiv 026062; doi: https://doi.org/10.1101/026062
[2] Bolstad BM (2016): ***preprocessCore: A collection of pre-processing functions***. R package version 1.36.0, https://github.com/bmbolstad/preprocessCore
[3] Emmert-Streib F, Glazko GV, Altay G, de Matos Simoes R: **Statistical inference and reverse engineering of gene regulatory networks from observational expression data.** *Front Genet* 2012, **3:**8.
